# Prophage induction, but not production of phage particles, is required for lethal disease in a microbiome-replete murine model of enterohemorrhagic *E. coli* infection

**DOI:** 10.1101/348706

**Authors:** Sowmya Balasubramanian, Marcia S. Osburne, Haley BrinJones, Albert K. Tai, John M. Leong

## Abstract

Enterohemorrhagic *Escherichia coli* (EHEC) colonize intestinal epithelium by generating characteristic attaching and effacing (AE) lesions. They are lysogenized by prophage that encode Shiga toxin 2 (Stx2), which is responsible for severe clinical manifestations. As a lysogen, prophage genes leading to lytic growth and *stx2* expression are repressed, whereas induction of the bacterial SOS response in response to DNA damage leads to lytic phage growth and Stx2 production both *in vitro* and in germ-free or streptomycin-treated mice.

Some commensal bacteria diminish prophage induction and concomitant Stx2 production *in vitro*, whereas it has been proposed that phage-susceptible commensals may amplify Stx2 production by facilitating successive cycles of infection *in vivo*. We tested the role of phage induction in both Stx production and lethal disease in microbiome-replete mice, using our mouse model encompassing the murine pathogen *Citrobacterrodentium* lysogenized with the Stx2-encoding phage Φ*stx_2dact_*. This strain generates EHEC-like AE lesions on the murine intestine and causes lethal Stx-mediated disease. We found that lethal mouse infection did not require that Φ*stx_2dact_* infect or lysogenize commensal bacteria. In addition, we detected circularized phage genomes, potentially in the early stage of replication, in feces of infected mice, confirming that prophage induction occurs during infection of microbiota-replete mice. Further, *C. rodentium* (Φ*stx_2dact_*) mutants that do not respond to DNA damage or express *stx* produced neither high levels of Stx2 *in vitro* or lethal infection *in vivo*, confirming that SOS induction and concomitant expression of phage-encoded *stx* genes are required for disease. In contrast, *C. rodentium* (Φ*stx_2dact_*) mutants incapable of prophage genome excision or of packaging phage genomes retained the ability to produce Stx *in vitro*, as well as to cause lethal disease in mice. Thus, in a microbiome-replete EHEC infection model, lytic induction of Stx-encoding prophage is essential for lethal disease, but actual phage production is not.

**Author summary:** Enterohemorrhagic *Escherichia coli* (EHEC), a food-borne pathogen that produces Shiga toxin, is associated with serious disease outbreaks worldwide, including over 390 food poisoning outbreaks in the U.S. in the last two decades. Humans acquire EHEC by ingesting contaminated food or water, or through contact with animals or their environment. Infection and toxin production may result in localized hemorrhagic colitis, but may progress to life-threatening systemic hemolytic uremic syndrome (HUS), the leading cause of kidney failure in children. Treatment for EHEC or HUS remains elusive, as antibiotics have been shown to exacerbate disease.

Shiga toxin genes reside on a dormant bacterial virus present in the EHEC genome, but are expressed when the virus is induced to leave its dormant state and begin to replicate. Extensive virus replication has been thought necessary to produce sufficient toxin to cause disease.

Using viral and bacterial mutants in our EHEC disease mouse model, we showed that whereas an inducing signal needed to begin viral replication was essential for lethal disease, virus production was not: sufficient Shiga toxin was produced to cause lethal mouse disease, even without viral replication. Future analyses of EHEC-infected human samples will determine whether this same phenomenon applies, potentially directing intervention strategies.

## Introduction

Shiga toxin-producing *Escherichia coli* (STEC) is a food-borne zoonotic agent associated with worldwide disease outbreaks that pose a serious public health concern. Enterohemorrhagic *Escherichia coli* (EHEC), a subset of STEC harboring specific virulence factors that promote a specific mode of colonization of the intestinal epithelium (see below), is acquired by humans by ingestion of contaminated food or water, or through contact with animals or their environment. EHEC serotype O157:H7 is a major source of *E. coli* food poisoning in the United States, accounting for more than 390 outbreaks in the last two decades[1–5]. EHEC infection usually presents as localized hemorrhagic colitis, and may progress to the life-threatening systemic hemolytic uremic syndrome (HUS), characterized by the triad of hemolytic anemia, thrombocytopenia, and renal failure [5, 6]. HUS is the leading cause of renal failure in children [7].

EHEC, along with enteropathogenic *E. coli* and *Citrobacter rodentium* belong to the family of bacteria known as attaching and effacing (AE) pathogens that are capable of forming pedestal-like structures beneath bound bacteria by triggering localized actin assembly [8–10]. While this ability of EHEC leads to colonization of the large intestine, production of prophage-encoded Shiga toxin (Stx) promotes intestinal damage resulting in hemorrhagic colitis [11–17]. Shiga toxin may further translocate across the colonic epithelium into the bloodstream, leading to systemic disease. Distal tissue sites, including the kidney, express high levels of the Shiga toxin-binding globotriosylceramide (Gb3) receptor, potentially leading to HUS [14, 15, 18–21].

Genes encoding EHEC Shiga toxin are typically encoded in the late gene transcription region of integrated lambdoid prophages [22, 23] and their expression is thus predicted to be temporally controlled by phage regulons [24–27]. Early studies showed that high levels of Stx production and release from the bacterium *in vitro* required prophage induction, i.e., the mechanism by which quiescent prophages of lysogenic bacteria are induced to replicate intracellularly and released as phage particles by host cell lysis [27, 28]. Lambdoid phage inducers are most commonly agents that damage DNA or interfere with DNA synthesis, such as ultraviolet light or mitomycin C. These inducing stimuli trigger activation of the bacterial RecA protein, ultimately leading to the cleavage of the prophage major repressor protein, CI, allowing expression of phage early and middle genes. Late gene transcription, which requires the Q antiterminator, results in the expression of many virion structural genes and of endolytic functions S and R, which lyse the bacterium and release progeny phage [29]. Other signaling pathways involving quorum sensing or stress response have also been implicated in lysogenic induction [30, 31].

Unfortunately, antibiotics commonly used to treat diarrheal diseases in children and adults are known to induce the SOS response‥ Trimethoprim-sulfamethoxazole and ciprofloxacin have been shown to enhance Stx production *in vitro* [32–34], and antibiotic treatment of EHEC-infected individuals is associated with an increased risk of HUS [35]. Hence, antibiotics are contraindicated for EHEC infection and current treatment is limited to supportive measures [36].

A more detailed understanding of the role of prophage induction and Stx production and disease has been pursued in animal models of EHEC infection. Although some strains of conventional mice can be transiently colonized with EHEC, colonization is not robust and typically diminishes over the course of a week [13, 37], necessitating use of streptomycin-treated [16] or germ-free mice [38, 39] to investigate disease manifestations that require efficient, longer term intestinal colonization. In streptomycin-treated mice colonized with EHEC, administration of ciprofloxacin, a known SOS inducer, induces the Stx prophage lytic cycle, leading to increased Stx production in mouse intestines and to Stx-mediated lethality [40]. Conversely, an EHEC strain encoding a mutant CI repressor incapable of inactivation by the SOS response was also incapable of causing disease in germ-free mice [41].

A potential limitation of the antibiotic-treated or germ-free mouse infection models is the disruption or absence, respectively, of microbiota, with concomitant alterations in immune and physiological function [42]. For example, a laboratory-adapted *E. coli* strain that lacks the colonization factors of commensal or pathogenic *E. coli* is capable of stably colonizing streptomycin-treated mice [43], and, when overproducing Stx2, is capable of causing lethal infection in antibiotic-treated mice [17]. Further, as up to 10% of human gut commensal *E. coli* were found to be susceptible to lysogenic infection by Stx phages *in vitro* [44], and it has been postulated that commensals may play an amplifying role in EHEC disease by fostering successive rounds of lytic phage growth [44–47]. Finally, gut microbiota may also directly influence expression of *stx* genes. For example, whereas a genetic sensor of phage induction suggests that the luminal environment of the germ-free mouse intestine harbors a prophage-inducing stimulus [41], several commensal bacteria have been shown to inhibit prophage induction and/or Stx production *in vitro* [48–50]. Alternatively, colicinogenic bacteria produce DNAse colicins that may trigger the SOS response, increasing Stx production [51].

Our laboratory previously developed a murine model for EHEC using the murine AE pathogen *C. rodentium* [52, 53], which efficiently colonizes conventionally raised mice and allows the study of infection in mice with intact microbiota. The infecting *C. rodentium* is lysogenized with *E. coli* Stx2-producing phage Φ1720a-02 [52, 54] encoding Stx variant Stx2_*dact*_, which is particularly potent in mice [55, 56]. Infection of C57BL/6 mice with *C. rodentium* (Φ1720a-02), (herein referred to as *C. rodentium* (Φ*stx_2dact_*)), produces many of the features of human EHEC infection, including colitis, renal damage, weight loss, and potential lethality, in an Stx2_dact_-dependent manner [52].

We have been interested in phage, bacterial, and host factors that lead to lethal EHEC infection. In the current study, we found that *C. rodentium* (*stx_2dact_*) strains lacking RecA, which is required for induction of an SOS response, or phage *Q* protein, which is required for efficient transcription of the late phage genes, did not produce high levels of Stx *in vitro* or cause lethal disease in mice. In contrast, mutants defective in prophage excision, phage assembly, or phage-induced bacterial lysis retained the ability to both produce Stx upon prophage induction *in vitro* and to cause lethal disease. Excised phage genomes, potentially undergoing DNA replication leading to phage production or representing packaged phage, were detected, albeit at low levels, in fecal samples of mice infected with wild type *C. rodentium* (φStx), but not in mice infected with excision-defective *C. rodentium* (φStx). Thus, in a microbiome-replete EHEC infection model, lytic induction of Stx-encoding prophage, but not actual production of viable phage particles, is essential for Stx production and lethal disease.

## Results

### Gene Map and Features of Φ*stx_2dact_* prophage

Lambdoid phage Φ1720a-02 was originally isolated from EC1720a-02, a STEC strain found in packaged ground beef [54]. To create the novel *C. rodentium-mediated* mouse model of EHEC infection, *C. rodentium* DBS100 (also known as *C. rodentium* strain ICC 168 (GenBank accession number NC_013716.1)), was lysogenized with phage Φ1720a-02 marked with a chloramphenicol (*cam*)-resistance cassette inserted into the phage *Rz* gene, creating lysogen DBS770 [52, 53]. A second lysogen, DBS771, encodes the chloramphenicol-marked prophage with an additional kanamycin (*kan*)-resistance cassette inserted into the prophage *stx2A* gene. Thus, in contrast to strain DBS770, DBS771 is unable to produce Shiga toxin or mediate lethal infection in mice. For simplicity, and for clarity with regard to phage mutations, phage Φ1720a-02, and strains DBS770 and DBS77′ will herein be referred to as Φ*stx_2dact_, C. rodentium* (Φ*stx_2dact_)*, and *C. rodentium* (ΦΔ*stx_2dact_::kan^R^*), respectively (Table 1).

**Table 1.**
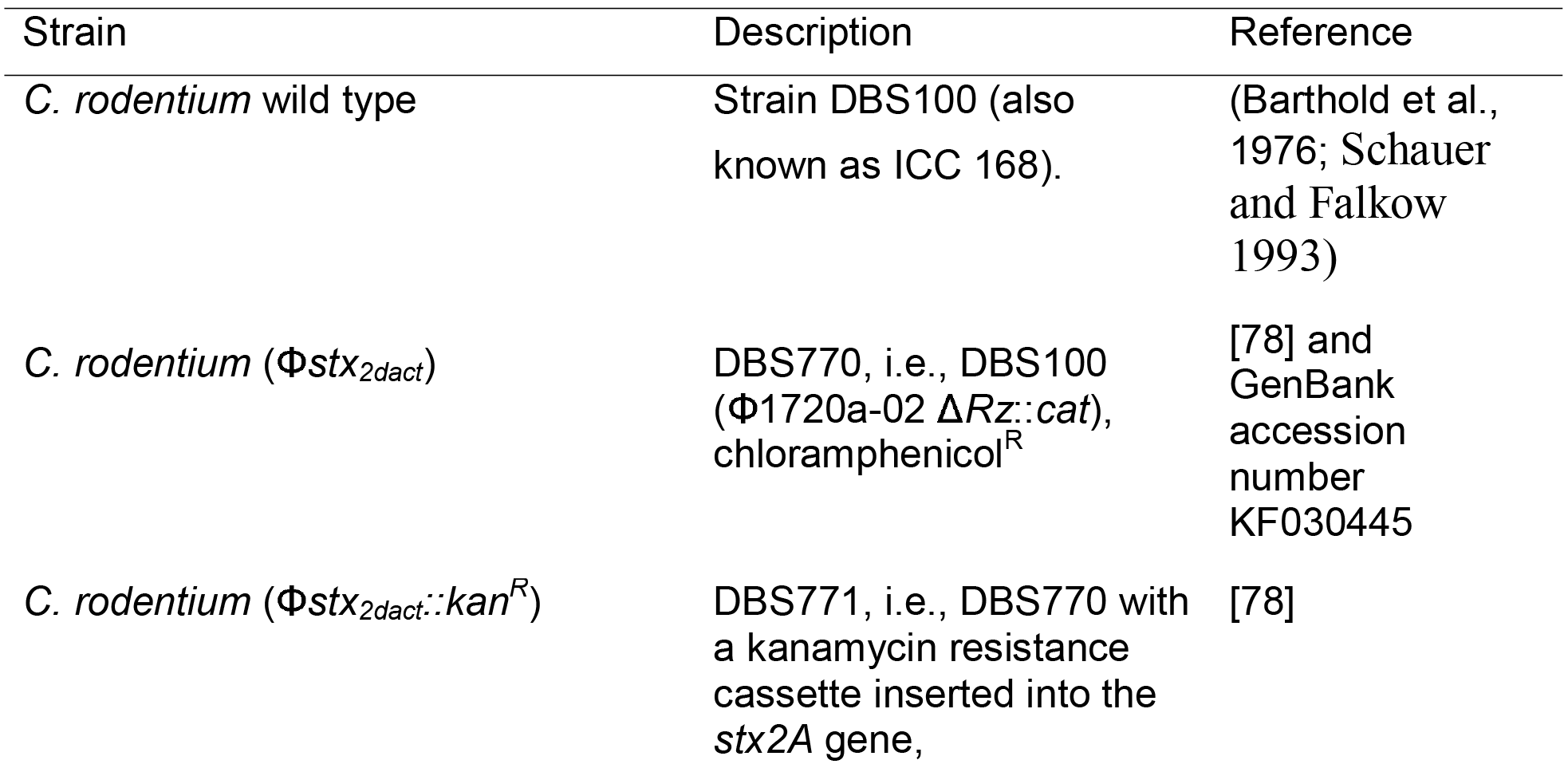
Bacterial strains and plasmids.

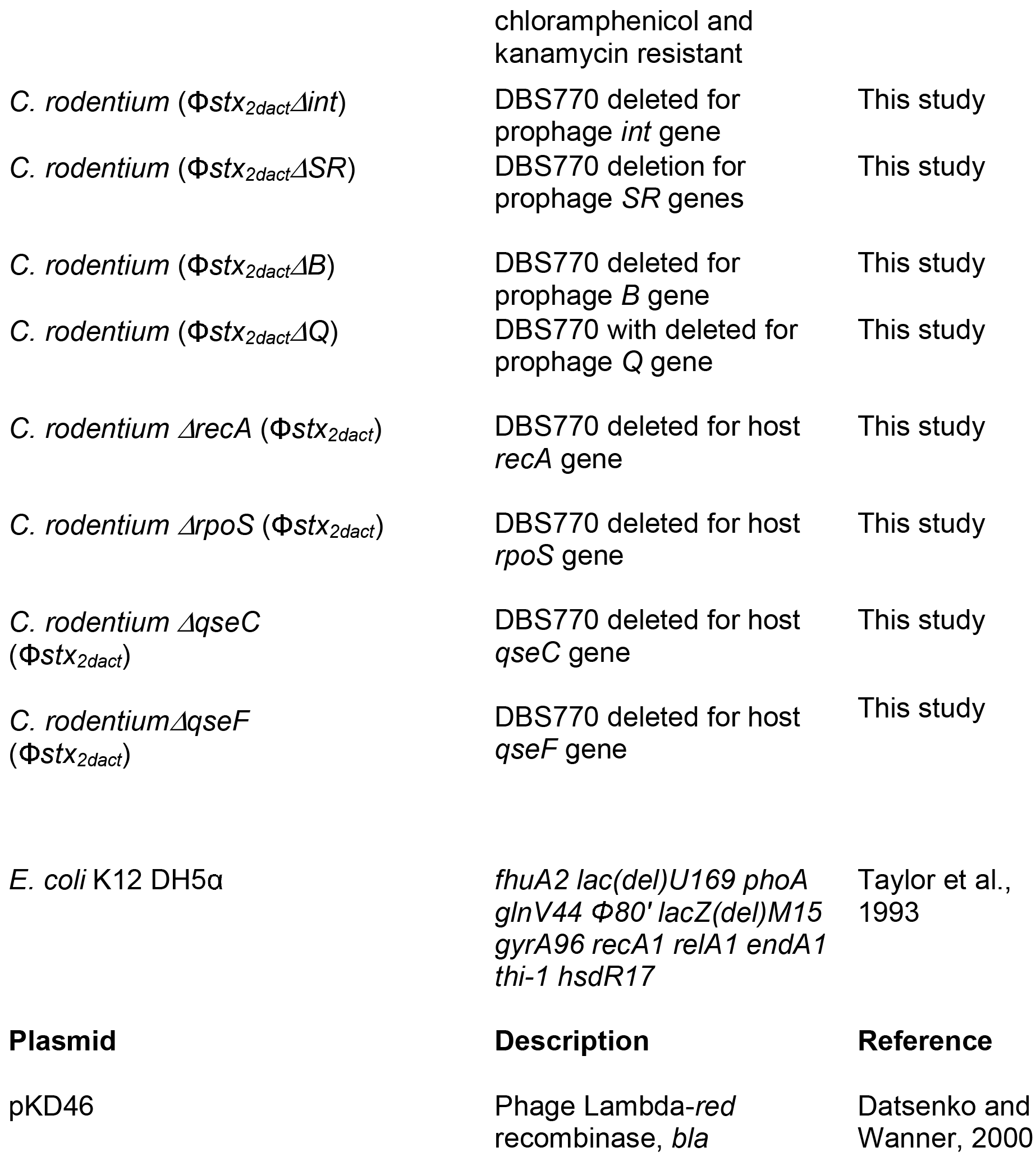

To identify phage genes critical for lethal mouse infection, we sought to inactivate specific prophage genes and then assess their resulting phenotypes in the *C. rodentium* mouse model. As a first step, we sequenced the parental strain DBS100 and the genomes of *C. rodentium* (Φ*stx_2dact_)* and *C. rodentium* (Φ*stx_2dact_::kan^R^*), revealing that the three genomes were identical except for the prophages present in *C. rodentium* (Φ*stx_2dac_t)* and *C. rodentium* (Φ*stx_2dact_::kan^R^*) (data not shown). We utilized these sequences to annotate the entire Φ*stx_2dact_*prophage (GenBank accession number KF030445.1) present in strains *C. rodentium* (Φ*stx_2dact_*) and *C. rodentium* (Φ*stx_2dact_*::Kan^R^), as diagrammed in Fig. S1. As is typical of Stx phages, the prophage sequence revealed a lambdoid phage with a mosaic gene organization, but nevertheless syntenic to varying degrees with other lambdoid phages [65].

Of note, as is the case for another lambdoid phage (*E. coli* phage mEP460, GenBank accession number JQ182728), the orientation of the *Φstx_2dact_* central regulatory region encoding CI repressor, other regulators such as N, Q and Cro, and the lytic promoters P_L_ and P_R_, is inverted with respect to the canonical map of phage lambda [66]. Although strains *C. rodentium* (Φ*stx_2dact_*) and *C. rodentium* (Φ*stx_2dact_::kan^R^*) were lysogenized independently, in both strains the prophage was integrated at the same location, i.e. 100 bp into the coding sequence of *dusA*, which encodes tRNA-dihydroxyuridine synthase A. A recent study [67] revealed that known integrase genes, at least half of which belong to prophages, were found adjacent to the host *dusA* gene in over 200 bacterial species. Furthermore, a 21 base pair motif found at the prophage-host DNA junctions in many bacteria was present at the prophage junctions, *attL* and *attR*, of *C. rodentium* (Φ*stx_2dact_*) and *C. rodentium* (ΦΔ*stx_2dact_::kan^R^*), as well as at the presumed *attB* phage insertion site in the parental *C. rodentium dusA* gene (Fig. 1). A seven-base segment within this 21-base sequence is completely conserved between *attL, attR*, and *attB* and likely represents the ‘core’ sequence within which recombination occurs during integration or excision (Fig. 1, bolded sequence; [68]). Finally, although the Φ*stx_2dact_* and ΦΔ*stx_2dact_::kan^R^* prophages interrupt the *dusA* gene, they encode a 184 bp ORF (designated “Φ*dusA’”* in Fig. 1) that is in frame with the 3’ 937 nucleotides (positions 101 to 1038) of *dusA* that could serve to foster the production of a protein containing the C-terminal 312 amino acids of the canonical DusA.

**Fig. 1.**
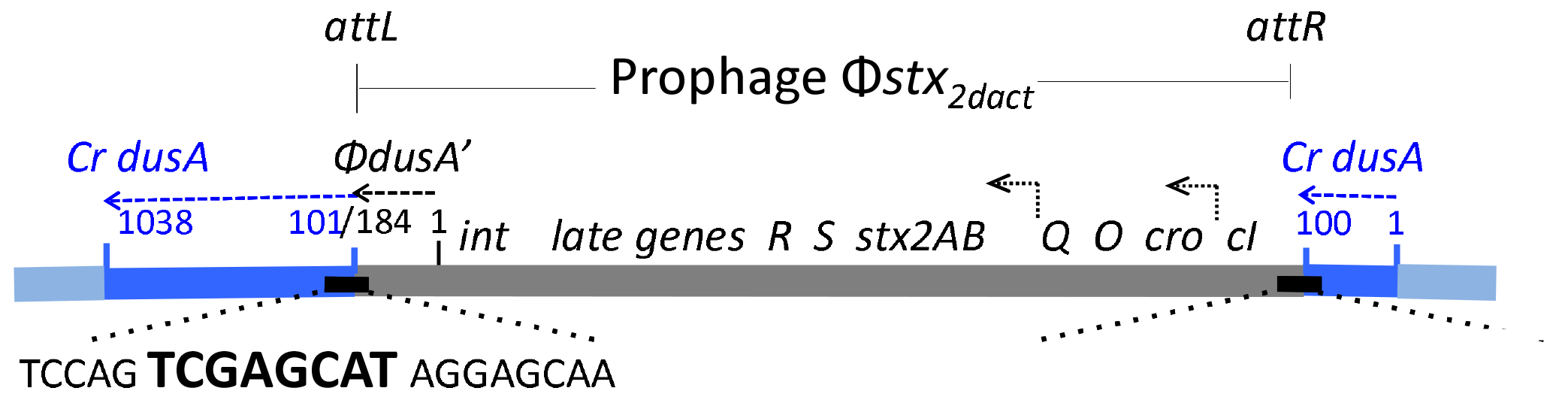
Prophage *stx_2dact_*in *C. rodentium* (*stx_2dact_*). The Φ*stx_2dact_* prophage (gray), flanked by *attL* and *attR* upon insertion into *C. rodentium dusA* sequence (blue, “*Cr dusA*”), was determined by whole genome sequencing of *C. rodentium (ΦΔstx_2dact_::kan^R^*). The 3’ end of the prophage (nucleotides 1-184) encodes the N-terminal 61 residues of “*ΦdusA*,”’ in the same reading frame as the 3’ end (nucleotides 101-1038) of the *C. rodentium dusA* gene (“*Cr dusA”*). Bent arrows indicate direction of transcription of Q, *stx*, and phage late genes. Depicted are *attL* and *attR* sequence motifs, characteristic of other prophages inserted within the host *dusA* gene ([67]). Within this sequence, a bolded seven-base “core” sequence, perfectly conserved in *attL* and *attR*, as well as in the Φ*stx_2dact_ attP* sequence and the parental *attB* sequence in *C. rodentium dusA* (not shown), is the cross-over site for phage integration and excision.

A prior analysis of the host *C. rodentium* DBS100 genome sequence revealed the presence of 10 additional partial and intact prophages distributed around the genome [69]. Using NCBI BLAST nucleotide analysis, we found only 2 regions of homology between Φ*stx2dactand* these prophages: one resident prophage (integrated at *C. rodentium* genome bp 2764517 – 2766051) encoded partial (70%) homology to the Φ*stx_2dact_ c*I repressor gene, and another resident prophage (integrated at *C. rodentium* genome bp 2097787-2098157) showed 79% homology to a Φ*stx_2dact_* gene encoding a hypothetical protein (data not shown). No other significant homology between Φ*stx_2dact_* and the resident prophages was detected.

### Survey of prophage integration (*attP*) sites during murine infection reveals Φ*stx_2dact_* prophage excision from *C. rodentium* (Φ*stx_2dact_*), but no secondary lysogeny of commensal bacteria

In the course of EHEC infection of streptomycin-treated mice, Stx phage could be induced by antibiotic treatment to lysogenize other *E coli* strains in the intraluminal environment [40], (also see [70]). It has been postulated that successive cycles of infection of non-pathogenic commensal *E. coli* could amplify Stx production and exacerbate disease [38, 44, 45, 47]. We first addressed this question by testing whether lysogeny of commensal bacteria by phage Φ*stx_2dac_t* was detectable following oral *C. rodentium (Φstx_2dact_*) infection of mice. Mice orally gavaged with *C. rodentium* (O*stx_2dact_*) normally exhibit weight loss and lethal disease ([52], not shown), typically succumbing to disease after day 7 post-infection. DNA was extracted from fecal samples of a group of five mice at days 1 and 6 post infection. The DNA samples were used as a template to generate a library of sequences encompassing the sequence downstream of *attL* (specifically, spanning the region from the phage *int* gene, through Φ*dusA* and into the adjacent host sequence; see Fig. 1). This strategy is a modification of that used for *Tn*-seq library analysis ([62], Materials and Methods).

Although we were unable to obtain detectable amplified DNA from fecal samples produced on day 1 post-infection, consistent with the low titer of *C. rodentium* (Φ*stx_2dact_*) in the stool at this early time point, the day 6 post-infection sample yielded a DNA library, which was subjected to massively parallel sequencing to identify the origin of the host DNA into which the prophage was integrated. Of the total 17,868,095 sequences generated, all but 725,997 (4.06%) were of sufficient quality to analyze (Table 2; see Materials and Methods). Of the readable sequences, 99.56% showed homology to *C. rodentium (Φstx_2dact_*), i.e. included *C. rodentium* (Φ*stx2dact*) *attL* and the adjacent *C. rodentium dusA* gene sequence, indicating prophage integration into the original *C. rodentium* strain. For the remaining 0.44% of sequences, the *C. rodentium dusA* sequences adjacent to the *attL* core sequence were found to be replaced by phage-specific sequences of *attR* at the other end of the prophage. Hence, these 0.44% of sequences encode *attP*, likely regenerated by recombination of *attL* and *attR* and probably reflecting excised circular phage genomes generated following induction of the *C. rodentium (Φstx_2dact_*) lysogen. Thus, our analysis indicates that *C. rodentium* (Φ*stx_2dact_*) undergoes lytic induction in the murine host, consistent with previous findings of EHEC infection in streptomycin-treated mice. Furthermore, of the more than 17 million sequences analyzed, none showed integration of the Φ*stx2dact* prophage into a
different site in *C. rodentium*, or into a different bacterial host. Thus, lysogeny of commensal bacteria by Φ*stx_2dact_* is not a common event in this model.

**Table 2.**
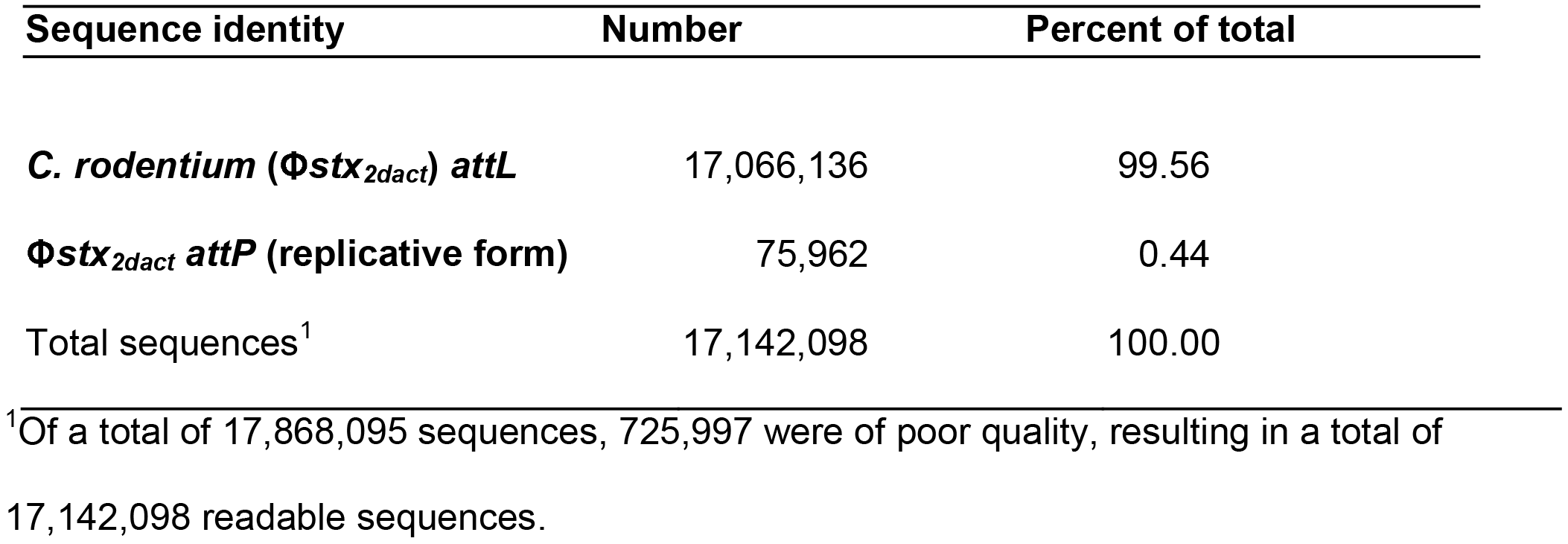
Comprehensive survey of prophage attachment (integration) sites reveals prophage excision but not secondary lysogeny of commensal bacteria during murine infection by *C. rodentium (Φstx_2dact_*)

| Sequence identity | Number | Percent of total |
| --- | --- | --- |
| *C. rodentium* ($\Phi stx_{2dact}$) *attL* | 17,066,136 | 99.56 |
| $\Phi stx_{2dact}$ *attP* (replicative form) | 75,962 | 0.44 |
| Total sequences<sup>1</sup> | 17,142,098 | 100.00 |

<sup>1</sup>Of a total of 17,868,095 sequences, 725,997 were of poor quality, resulting in a total of 17,142,098 readable sequences.

### *C. rodentium* RecA and Φ*stx*_*2dact*_ proteins integrase, Q, endolysins, and portal protein are required for efficient phage production and release *in vitro*

Prophage induction of lambdoid phages is often initiated by DNA damage, in which SOS pathway activation leads to RecA-promoted autocleavage of CI repressor, followed by :ranscription of early genes from the from P_L_ and P_R_ promoters. Subsequent temporally programmed transcription of the prophage genome results in the production of delayed early (middle) proteins such as Int (integrase), essential for prophage integration and excision, and antiterminator protein Q. Production of Q in turn mediates the transcription of late genes, ncluding portal protein gene *B*, responsible for translocation of phage DNA into the virion protein capsid, and lysis genes *S* and *R*, encoding endolysins that disrupt the bacterial plasma membrane causing release of intact phage progeny (for a review, see Gottesman and Weisberg [66]). Late genes in EHEC phages also encompass *stx*.

To uncover the roles of specific phage and bacterial functions in EHEC disease, we used lambda *red* recombination (Materials and Methods) to construct *C. rodentium* (Φ*stx_2dact_*) strains defective for prophage genes *SR, int, B*, or *Q*, and the host gene *recA*, which is well documented to be central to the SOS response and lytic induction. In addition, we inactivated three other genes that have been implicated as having more subtle roles in the lytic induction of Shiga toxin-encoding phage [30, 31]: *rpoS*, which controls the bacterial stress response, and *qseC* and *qseF*, which control quorum sensing pathways (Materials and Methods, Table 1). None of the mutants displayed a growth defect upon *in vitro* culture in rich broth (Fig. S2 and data not shown).

We then tested *C. rodentium* (Φ*stx_2dact_*) and several of the mutant derivatives predicted to have dramatic effects on phage production for the ability to generate Φ*stx_2dact_* following SOS induction. Pilot experiments revealed that Φ*stx_2dact_* plaques were not detectable on any of numerous indicator strains (not shown), so phage production was measured by qPCR [71] using primers flanking the phage *attP* site (Table 3). As only excised phage have a reconstituted *attP* site [66], these primers only amplify product from unintegrated phage genomes.

**Table 3.**
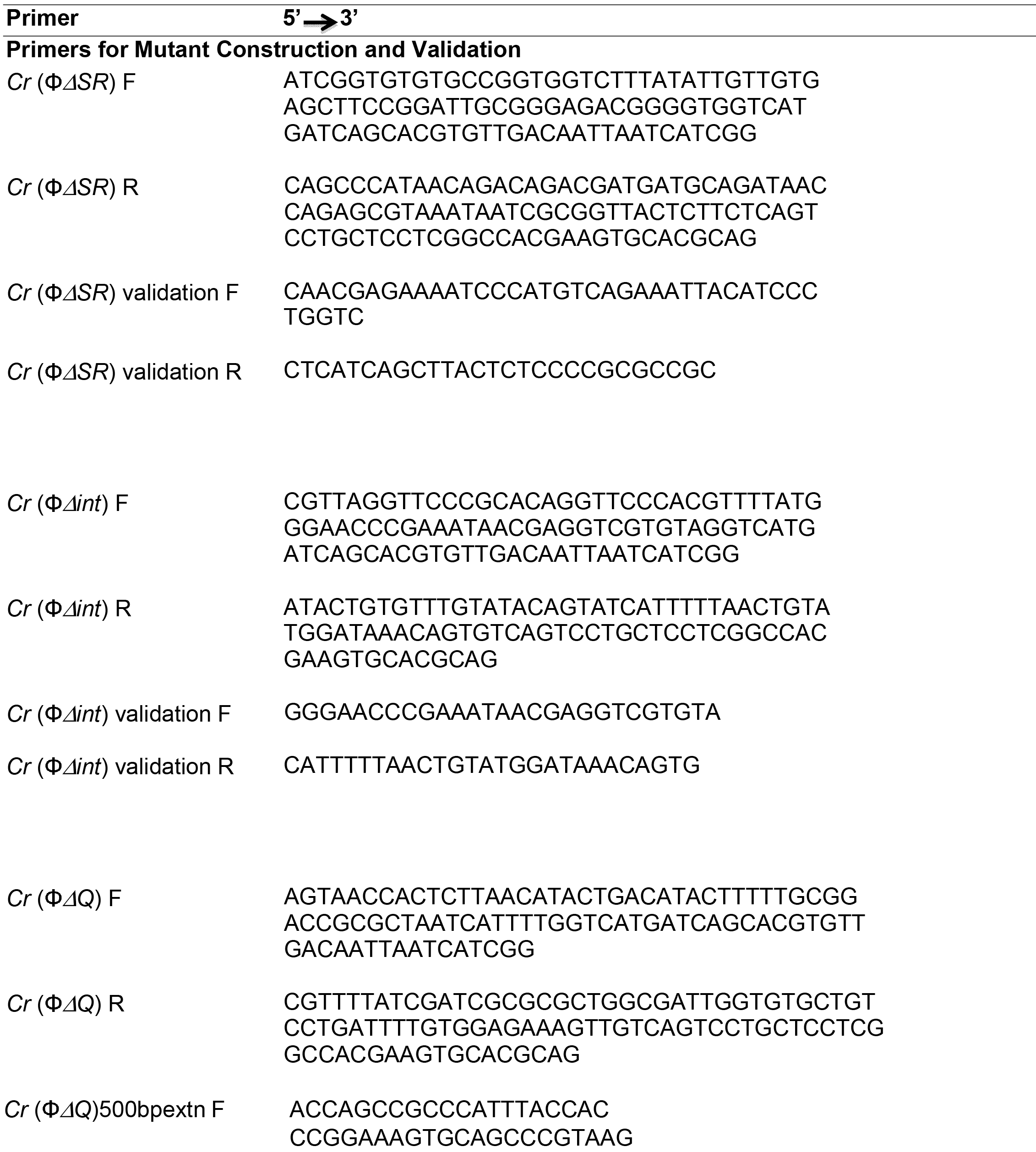
Primers used in this study.

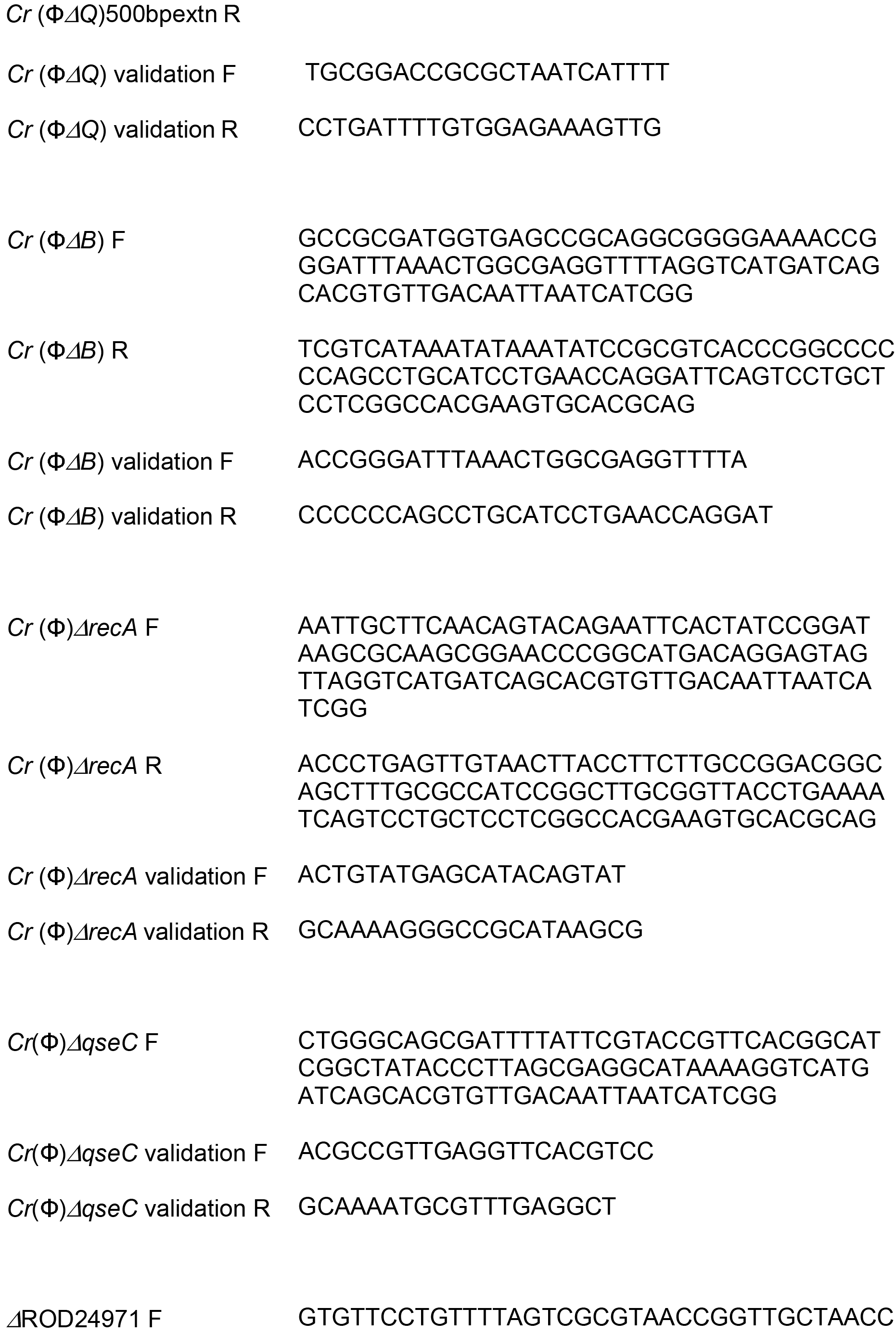

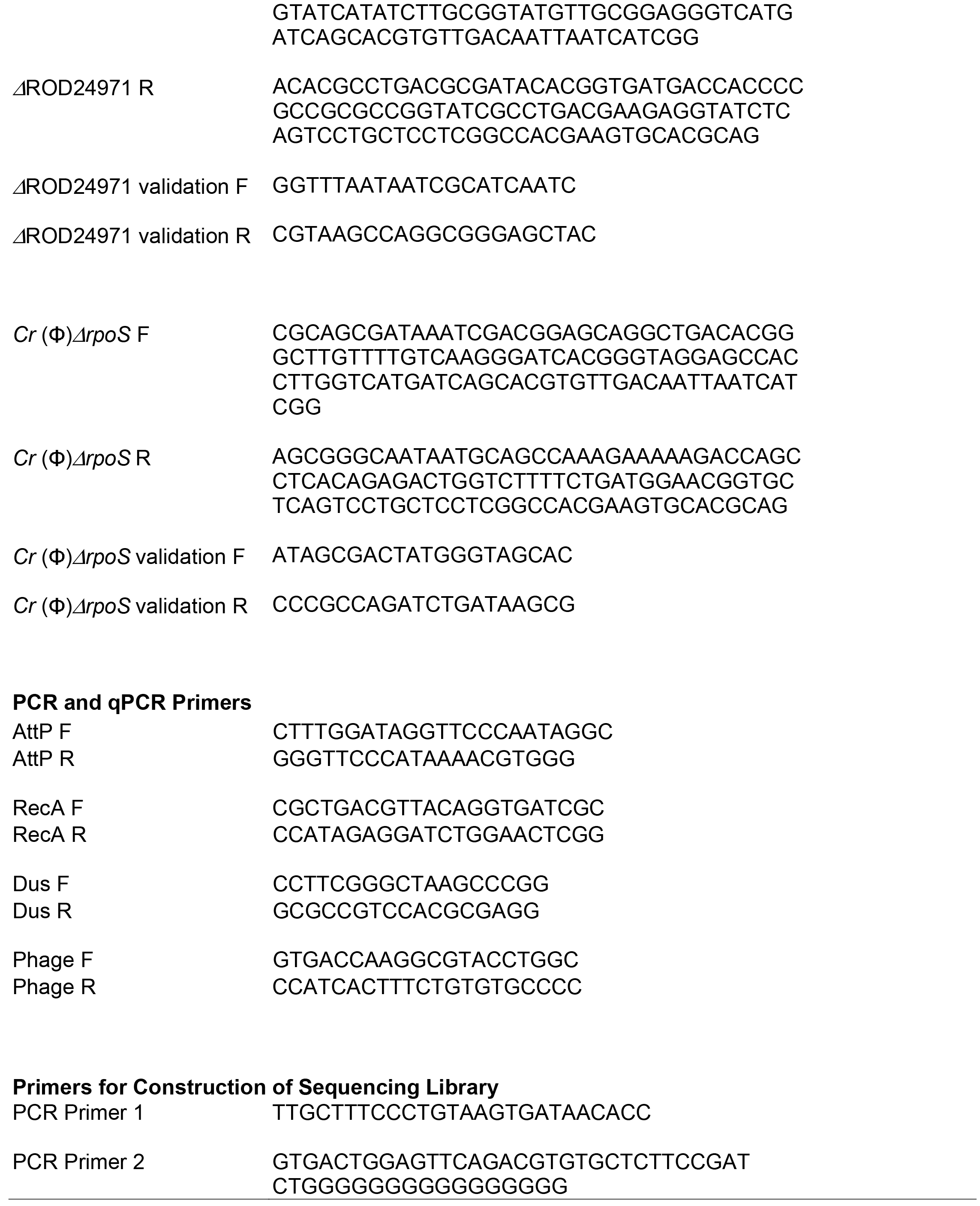

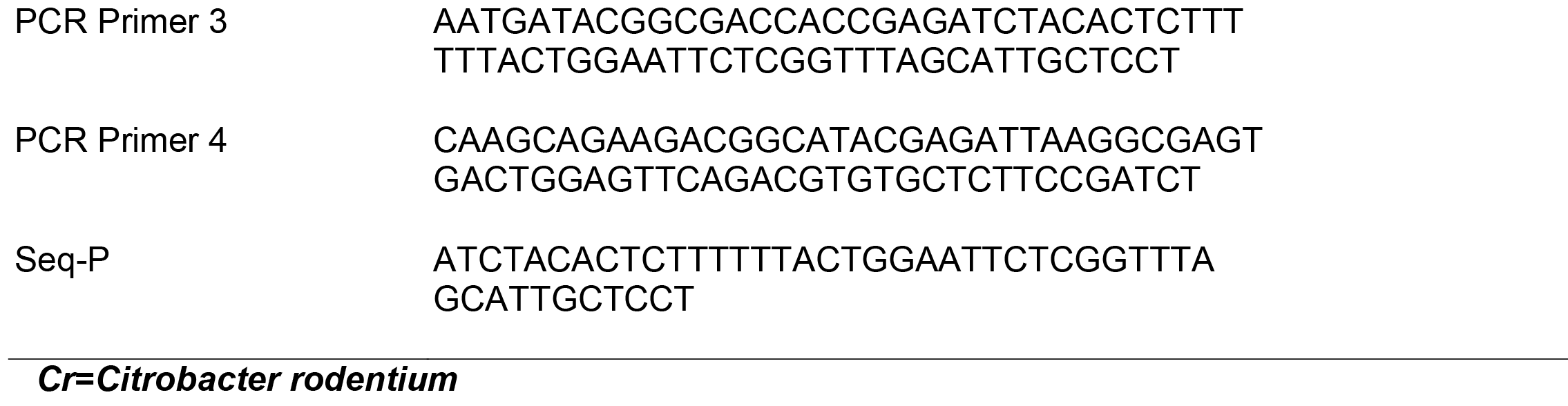

In supernatants of LB cultures at mid-logarithmic growth phase, designated as t=0h, we detected 1.3⨯10^9^ - 3.8⨯10^9^ *attP* copies (phage genomes) per ml (Table 4 legend). The midlog cultures of wild type *C. rodentium* (Φ*stx_2dact_*) contained only approximately 10^8^ viable bacteria per ml, indicating that significant spontaneous prophage induction had occurred during the 2-4 hours of culture of this strain prior to entering mid-log growth. Four hours of further culture of *C. rodentium* (Φ*stx_2dact_*) (t=4h) resulted in a 3.2-fold increase in the concentration of Φ*stx_2dact_* in the supernatant compared to that at t=0h, consistent with continued spontaneous prophage induction (Table 4, “Relative *attP* production”, “- Mito C”). Prophage induction of the wild type lysogen with the SOS inducer mitomycin C led to a 234fold increase in relative *attP* production (Table 4, “+ Mito C”), a 73-fold increase above baseline levels. As predicted [72, 73], the generation of circular phage genomes required integrase; at all time points tested, *attP* copies were below the level of detection of 1⨯ 10^4^/ml in uninduced or mitomycin C-induced cultures of the *C. rodentium* (Φ*stx_2dact_*Δ*int*) mutant, which lacks Int recombinase (Table 4).

**Table 4.** *C. rodentium* RecA and Φ*stx_2dact_* proteins integrase, Q, endolysins, and portal protein are required for efficient phage production and release *in vitro*.

| Strain | Function deleted | Relative *attP* (phage) production |  |  |
| --- | --- | --- | --- | --- |
|  |  | - Mito C<sup>a</sup> | + Mito C<sup>b</sup> | +Mito C + DNase<sup>c</sup> |
| WT | None (WT) | 3.2 ($\pm 0.01$) | 234.6 ($\pm 24.4$) | 162.5 ($\pm 1.1$) |
| $\Delta\text{int}$ | Phage integrase | Not detected | Not detected | Not determined |
| $\Delta\text{recA}$ | Host RecA | 0.5 ($\pm 0.6$)* | 29.4 ($\pm 11.4$)* | Not determined |
| $\Delta\text{Q}$ | Phage late gene transcription anti-terminator | 0.5 ($\pm 0.3$)* | 6.0 ($\pm 0.3$)* | Not determined |
| $\Delta\text{SR}$ | Phage endolysin | 0.6 ($\pm 0.2$)* | 6.3 ($\pm 2.0$)* | Not determined |
| $\Delta\text{B}$ | Phage portal protein | 4.1 ($\pm 0.08$) | 208.7 ($\pm 17.2$) | 9.2 ($\pm 1.3$)* |

<sup>a</sup>Supernatants from mid-log (t=0h) cultures or parallel cultures grown for an additional 4 hours (t=4h) were analyzed for *attP* copies by qPCR. Shown are average values of t=4h/t=0h (+/- SEM) for each lysogen, derived from the values of three different dilutions of each supernatant (see Materials and Methods). For all lysogens except *C. rodentium* ( $\Phi\text{stx}_{2dact}\Delta\text{int}$ ), absolute numbers of *attP* molecules at t=0h ranged from $1.3 \times 10^9$ to $3.8 \times 10^9$ /ml. For *C. rodentium* ( $\Phi\text{stx}_{2dact}\Delta\text{int}$ ), *attP* copies were below the limit of detection, i.e., $<1 \times 10^4$ /ml.
<sup>b</sup>Supernatants from mid-log (t=0h) cultures, and parallel cultures subsequently exposed to 0.25 $\mu\text{g}/\text{ml}$ mitomycin C for 4 hours (t=4h) were analyzed for *attP* copies by qPCR and the ratios of the two values determined as above. For *C. rodentium* ( $\Phi\text{stx}_{2dact}\Delta\text{int}$ ), *attP* copies were below the limit of detection, i.e., $<1 \times 10^4$ /ml.

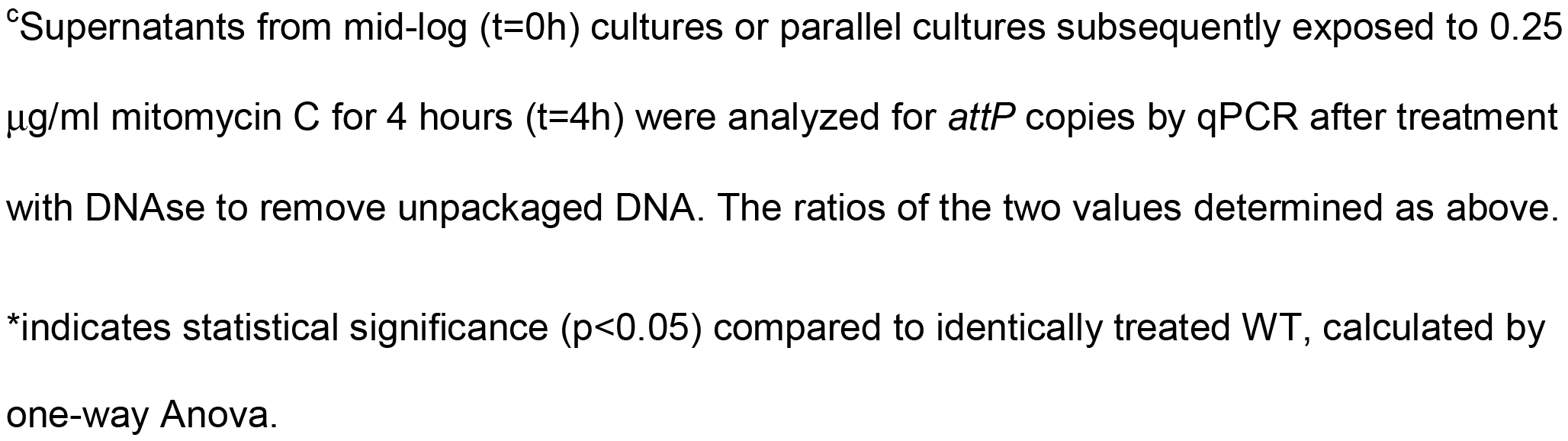

Host and phage functions contributed to the amount of phage production in both the absence and presence of inducer. In the absence of inducer, (Table 4, “- MitoC”) the concentration of *attP* copies in culture supernatants of *C. rodentium* Δ*recA* (Φ*stx_2dact_*), predicted to be defective for SOS induction, did not increase between t=0h and t=4h, with an average relative *attP* production of 0.5. Lysogens deficient in the antiterminator Q, required for late gene transcription, or deficient in the S and R endolysins, which promote the efficient release of phage from infected bacteria, were also deficient in relative *attP* production in the absence of inducer (Table 4). Finally, *C. rodentium* (Φ*stx_2dac_*Δ*B*), predicted to replicate but not package phage genomes, showed no defect in the production of *attP* copies in the culture supernatant in the absence of inducer, with relative phage production ratio of 4.1. However, as described below, DNAse sensitivity assays suggested that these *attP* sequences are likely not packaged into phage particles.

The mutants defective in baseline phage production were similarly defective in the titer of *attP* copies after induction with mitomycin C (Table 4, “+ Mito C”). Induction of *C. rodentium* Δ*recA* (Φ*stx_2dact_*) resulted in a relative *attP* production value of only 29, i.e. eight-fold lower than wild type. *C. rodentium* (Φ*stx_2dact_*Δ*Q*), and *C. rodentium* (Φ*stx_2dact_*Δ*SR*), each also demonstrated dramatically diminished *attP* copies in mitomycin C-induced culture supernatants, with relative *attP* production of approximately 6. Finally, *C. rodentium* (Φ*stx_2dact_*Δ*B*), generated wild type levels of phage genome copies, with a 209-fold increase in relative *attP* production. However, DNAse treatment of supernatants diminished this value more than 23-fold, whereas parallel treatment diminished the relative *attP* production by wild type *C. rodentium* (Φ*stx_2dact_*) less than 1.5-fold (Table 4, “+Mito C + DNAse”), consistent with a defect in packaging of Φ*stx_2dact_* genomes in the absence of the B portal protein.

### Proteins required for the SOS response and/or late gene transcription are essential for Stx2dact production

To determine which host or phage functions are required for production of Stx2dact *in vitro*, we measured Stx2dact in culture supernatants by ELISA [53]. To quantitate non-induced levels of Stx, and to provide ample time for toxin to accumulate, we grew triplicate cultures of the *C. rodentium* (Φ*stx_2dact_*) or the mutant derivatives described above for four hours (t=4h) beyond mid-log phase (defined as t=0h). Stx2dact was present in the culture supernatants of wild type *C. rodentium* (Φ*stx_2dact_*) at approximately 50 ng/ml/OD_600_ unit, consistent with previous measurements [53]. Prophage excision and phage production were not required for this basal level of Stx2dact: culture supernatants of *C. rodentium* (Φ*stx_2dact_*Δ*int*), which did not harbor detectable phage (Table 4), contained equivalent amounts of toxin (Fig. 2A, “Δ*int*”). Uninduced culture supertants of *C. rodentium* (Φ*stx_2dact_*Δ*SR*) contained levels of Stx2dact two-fold lower than (and statistically indistinguishable from) wild type, consistent with the moderately (5-fold) lower levels of phage found in cultures of wild type *C. rodentium* (Φ*stx_2dact_*) (Table 4, “- MitoC”). Supernatants of *C. rodentium* (Φ*stx_2dact_*Δ*B*), which contained *attP* DNA but relatively few packaged phage (Table 4), also produced levels of Stx2dact statistically indistinguishable from wild type. Finally, in contrast, *C. rodentium* Δ*recA* (Φ*stx_2dact_*), which is unable to mount an SOS response, and *C. rodentium* (Φstx_2dact_ΔQ), which cannot transcribe phage late genes, including *stx2dactA* and *stx2dactB*, were defective for basal levels of Stx2dact production (Fig. 2A, “Δ*recA”*, “ΔQ”).

**Fig 2.**
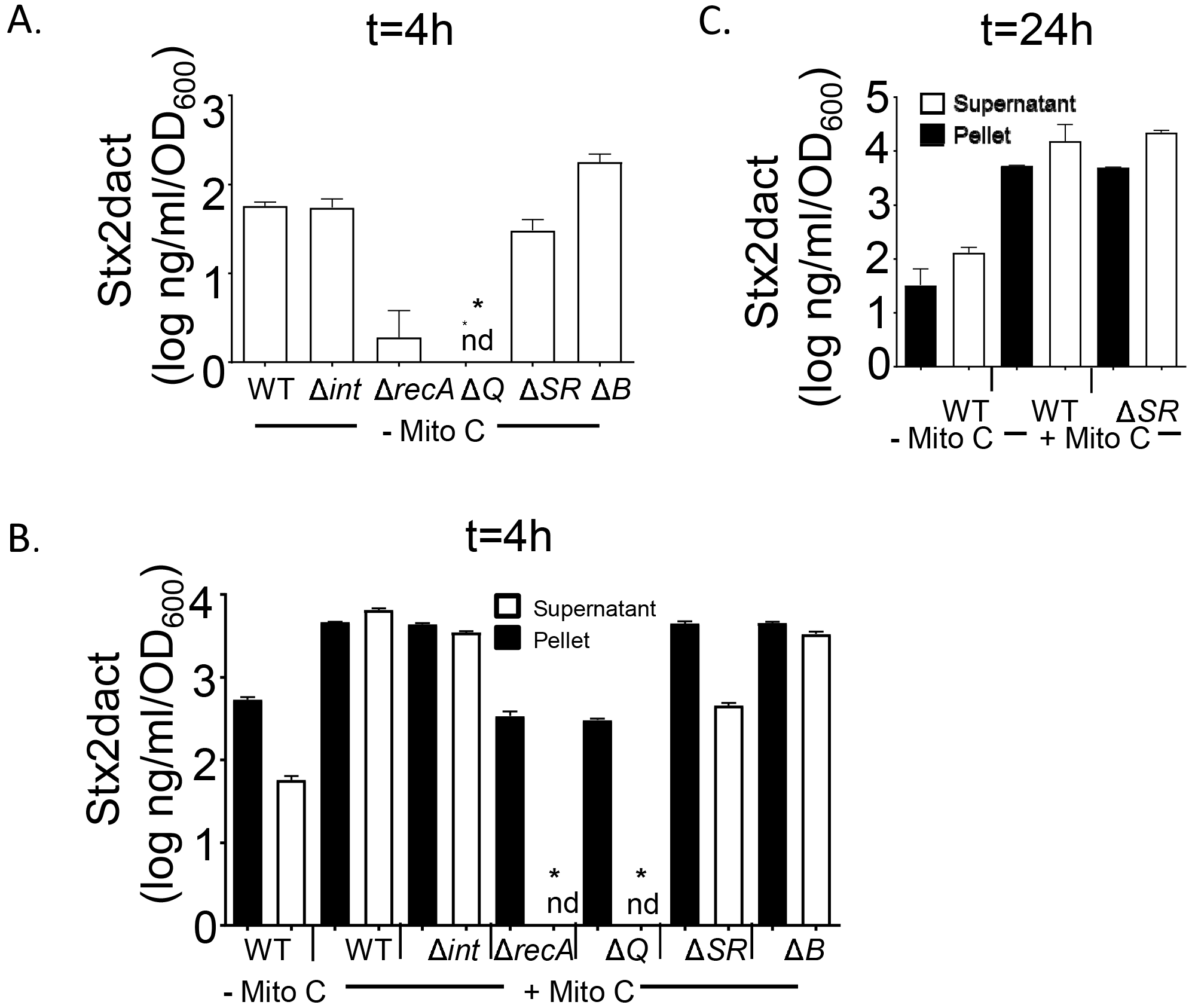
SOS responsiveness and lytic induction-dependent transcription of *stx* genes are required for wild type basal and induced levels of Stx2 production *in vitro*. **A.** The indicated lysogens were grown in the absence of mitomycin C until t=4h i.e., four hours after attaining approximately mid-log phase (which was designated as t=0h; “- Mito C“), and culture supernatants were subjected to capture ELISA to determine the basal level of Stx2 production (see Materials and Methods). Quantities are expressed relative to the specific OD_600_ at t=0h. B. The indicated lysogens were grown to mid-log phase (t=0h) and cultured for four more hours (t=4h) either in the absence (“- Mito C“) or presence of 0.25 `μ` g/ml mitomycin C (“+ Mito C“). Pellets (filled bars) or supernatants (open bars) were subjected to capture ELISA to determine the level of Stx2 production. Quantities are expressed relative to the specific OD_600_ at t=0h. **C.** Wild type *C. rodentium* (Φ*stx_2dact_*) and *C. rodentium* (Φ*stx_2dact_* ARS)) were grown to mid-log phase (designated as t=0h) and cultured for 16 more hours (t=16h) either in the absence (“- Mito C”) or presence of 0.25 `μ` g/ml mitomycin C (“+ Mito C”). Pellets (filled bars) or supernatants (open bars) were subjected to capture ELISA to determine the level of Stx2 production. Quantities are expressed relative to the specific OD_600_ at t=0h. For all panels, results are averages ± SEM of triplicate samples, and are a representative of at least two experiments involving independently derived mutants. Asterisks (*) indicate Stx level significantly (p <0.05) different from wild type *C. rodentium* (Ostx2dact) calculated using Kruskal-Wallis one-way analysis of variance followed by Dunn’s nonparametric comparison.

We also assessed Stx production by wild type *C. rodentium* (Φ*stx2dact*) and mutant derivatives after 4h of mitomycin C induction. Given that mitomycin C-induced Φ*stx_2dact_* functions may be involved in the release of toxin from the bacterial host [27], we assessed toxin in cell pellets and in culture supernatants separately. As previously observed [52], mitomycin C induction resulted in a more than 100-fold increase of Stx2dact in culture supernatants (Fig. 2B, “WT”). A nearly equivalent amount of toxin remained associated with the bacterial cell pellet, suggesting that under these conditions, a significant fraction of bacteria remained unlysed. Culture supernatants or cell pellets of the *C. rodentium ΔrpoS* (Φ*stx_2dact_*) mutant predicted to be defective in the bacterial stress response, or the *C. rodentium* Δ*qseC* (Φ*stx_2dact_*) mutant defective for quorum sensing, showed wild type levels of Stx2dact (Fig. S3), as did *C. rodentium* Δ*qseF* (Φ*stx_2dact_*) (data not shown), indicating that neither the bacterial stress response nor the QseC- or QseF-mediated quorum responses were required for toxin production. Culture supernatants of *C. rodentium* (Φ*stx_2dact_*Δ*int*) and *C. rodentium* (Φ*stx_2dac_*Δ*B*), which showed no defect in basal levels of toxin production (Fig. 2A), also contained amounts of cell-associated toxin and supernatant-associated Stx2dact indistinguishable from wild type (Fig. 2B, “Δ*int*” and “ Δ*B*”), despite the lack of prophage excision and/or phage production in these mutant strains. The ΔSR lysogen, defective for phage endolytic functions, produced wild type levels of cell-associated Stx2dact at 4 h postinduction, but supernatant-associated toxin was approximately ten-fold lower than wild type levels (Fig. 2B, “Δ *SR*”). This difference is consistent with a defect in bacterial lysis and Stx2dact release, but did not reach statistical significance. In addition, by 16 h post-induction of the Δ*SR* lysogen, Stx2dact was detected in supernatants at levels similar to that of the WT
strain (Fig. 2C), suggesting that any defect in R and S proteins results in a delay rather than an absolute block in toxin release. Finally, however, deficiency in the RecA or Q proteins was associated with a near-complete absence of Stx2dact in cell pellets or supernatants (Fig. 2B, “Δ*recA*” and “ Δ*Q*”), reinforcing the notion that these proteins, which are required for the SOS response and/or transcription of the *stx2_dact_* ([74] [66]) are essential for Stx2dact production.

### *C. rodentium* (Φ*stx_2dact_*) undergoes lytic induction during murine infection

Stx-encoding prophages undergo lytic induction during EHEC infection of germ-free or antibiotic-treated mice [40, 41, 70], and our comprehensive survey of prophage integration sites in fecal microbiota (Table 3) indicated that *C. rodentium* (Φ*stx_2dact_*) undergoes some degree of lytic induction during infection of conventional mice. To assess this induction further, we infected conventionally raised C57BL/6 mice with *C. rodentium* (Φ*stx2dact*) by oral gavage and measured fecal shedding of both the infecting strain, by plating for CFU, and Φ*stx_2dact_*, by quantitating *attP* (non-integrated phage) copies by qPCR. As previously observed, by day 3 post-infection, *C. rodentium* (Φ*stx_2dact_*) was detected in the stool at 8 ⨯ 10^7^ per gram, and reached 9 ⨯ 10^10^ per gram by day 6 post-infection ([64]; Fig. 3, “CFU of WT”). Further, murine infection by this strain was indeed associated with lytic induction, as excised phage genomes were detected in stool at all time points (Fig. 3, “Phage from WT”).

**Fig 3.**
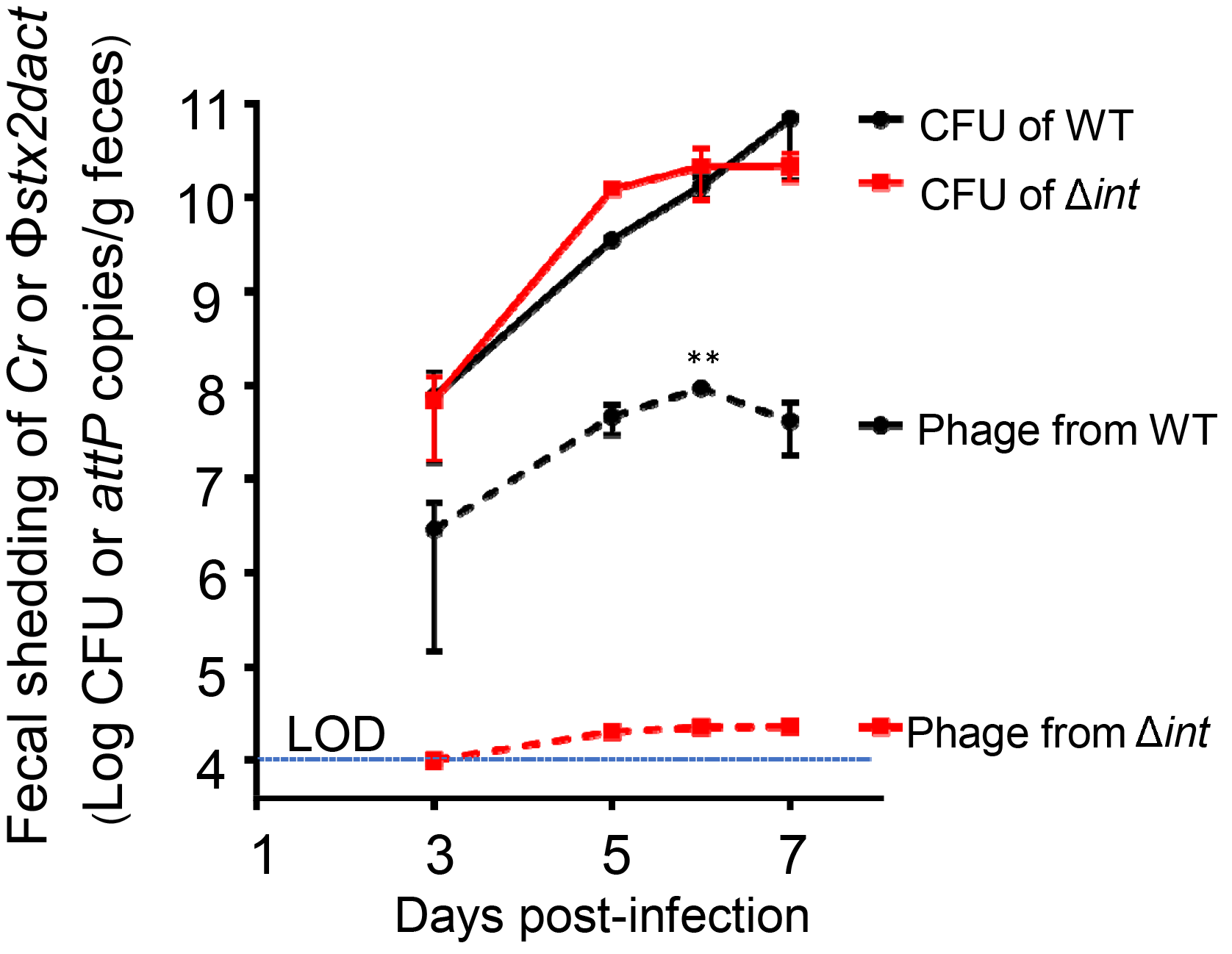
*C. rodentium (Φstx_2dact_*) undergoes lytic induction during murine infection. Eight-week old female C57BL/6 mice were infected by oral gavage with *C. rodentium* (Φ*stx_2dact_*) or *C. rodentium* (Φ*stx_adact_* Δ*int*). At the indicated time points, *attP* copies, reflecting excised prophages, and viable bacteria were determined by qPCR or plating for CFU, respectively (see Maaterials and Methods). Shown are averages ± SEM of 5 mice per group of a representative of two experiments. Level of detection of *attP* was 1 ⨯ 10^4^ copies/g feces. Asterisks (**) indicate significance differences (p <0.01) between the WT and *C. rodentium* (Φstx2_*dact*_ Δ*int*) calculated using 2-way ANOVA followed by Bonferroni post tests.

Interestingly, given the relatively high phage production by *C. rodentium* (Φ*stx_2dact_*) *in vitro*, the amount of phage detected in stool was quite low. At day 3 post-infection, 5 ⨯ 10^6^ *attP* copies were detected per gram of stool, a value 16-fold lower than the concentration of viable *C. rodentium* (Φ*stx_2dact_*) in stool at that time point. By day 6 post-infection, *attP* copies had increased to 5 ⨯ 10^7^ per gram of feces, but were approximately 600-fold lower than the fecal bacterial counts. *C. rodentium* Φ(*stx2dact*) thus undergoes lytic induction and growth in this murine model, although not to the degree seen *in vitro*.

### Lethal disease in mice correlates with the ability to produce Stx2dact but not with the ability to produce phage

To test the importance of SOS induction and phage functions on disease in our microbiota-replete model of infection, we infected C57BL/6 mice with *C. rodentium* (Φ*stx_2dact_*) and mutant derivatives by oral gavage. The wild type and all mutant lysogens colonized mice similarly (Fig. S4). *C. rodentium* Δ*recA* (Φ*stx2dact*) and *C. rodentium* (Φ*stx_2dact_*Δ*Q*), the two mutant lysogens that displayed dramatic defects in basal and mitomycin C-induced levels of Stx2dact *in vitro*, were the only ones incapable of causing sickness or death (Fig. 3, “Δ*recA*” and “ Δ*Q*”), supporting the hypothesis that induction of an SOS response and the subsequent expression of phage late genes, including *stx* genes, are required for Shiga toxin production during infection of a microbiota-replete host.

The RpoS-deficient and QseC-deficient *C. rodentium* (Φ*stx_2dact_*) mutants that are compromised in bacterial stress and quorum-sensing responses, respectively, retained the ability to cause weight loss and lethality with kinetics that were indistinguishable from that of WT *C. rodentium*(Φ*stx_2dact_*) (Fig. S5). Thus, although previous results indicated that some quorum sensing mutants display diminished virulence during infection by non-Stx-producing *C. rodentium* [75], our results are consistent with the the ability of these strains to produce wild type levels of Stx2 after SOS induction (Fig. S3). In addition, the lack of endolysins that appeared to somewhat delay release of Stx2dact into supernatants by *C. rodentium* (Φ*stx_2dact_*Δ*SR*) was not reflected by any delay in the kinetics of weight loss or lethality in infected mice (Fig. 3, “Δ*SR*”), consistent with the ability of this strain to produce wild type levels of Stx2dact upon extended culture *in vitro*.

Finally, the production of intact phage is not essential to disease in this model. *C. rodentium* (Φ*stx_2dact_*Δ*B*), which is unable to generate intact phage *in vitro*, and *C. rodentium* (Φ*stx_2dact_*Δ*int*), which can neither generate excised phage genomes *in vitro* or *in vivo*, both retained full virulence in this model. We conclude that in this microbiota-replete model of EHEC infection, disease progression correlates exclusively with the ability to produce Stx2, regardless of the lysogen’s ability to amplify the *stx2* gene by phage excision and genome amplification, or by the production of phage that are capable of secondary infection of commensal bacteria.

## Discussion

Commensal organisms have the potential to suppress or enhance phage induction and Stx production. Although a role for induction of *stx*-encoding prophages in the production of Stx and serious disease during animal infection has been well documented in antibiotic-treated and germ-free mice [40, 41, 70], we used a murine model of EHEC infection that features an intact microbiome.

To investigate phage functions required for *C. rodentium* (Φ*stx_2dact_*) to produce Stx and cause disease in conventional mice, we first characterized prophage genetic structure. Φ*stx_2dac_t* prophage was integrated into the *C. rodentium dusA* gene, an integration site utilized by prophages in over 200 bacterial species [67]. Although the orientation of the regulatory and late genes within the Φ*stx_2dact_* prophage is noncanonical with respect to *attL* and *attR* (with *int* adjacent to *attL*; Fig. 1), this orientation has been previously observed in at least one other lambdoid phage. In addition, Φ*stx_2dact_* genes encoding several key phage proteins were identified by homology, and their inactivation had the predicted effects on phage development and production (Table 4; [73]). For example, antiterminator Q and integrase were required for phage production, as measured by detection of *attP*, and portal protein B appeared to be required for packaging of phage DNA into DNAse-resistant virions.

Stx production *in vitro* by the prophage mutants, as well as by a host *recA* mutant, confirmed that prophage induction, i.e., the SOS-dependent process required to initiate a temporal program of phage gene expression that normally leads to phage lytic growth, is essential for high-level Stx2 production *in vitro*. Mitomycin C treatment of *C. rodentium* (Φ*stx_2dact_*) resulted in a greater than 100-fold increase in Stx2dact in culture supernatants, similar to the mitomycin C-mediated increase in Shiga toxin production by EHEC ([41]; Fig. 2). Two signaling pathways, mediated by RpoS and QseC, previously demonstrated to influence SOS induction of EHEC *in vitro*, had no effect on Stx2dact production by *C. rodentium* (Φ*stx_2dact_)*. In contrast, and as expected, RecA, required for mounting an SOS response, was necessary for this enhanced production of Stx2dact (Fig. 2). It was previously shown that inactivation of the EHEC prophage repressor CI, a key step in the SOS response, is required for the increase in EHEC Stx production upon mitomycin C induction *in vitro* [41].

Despite the previous observation that the increase in phage genome copy number plays the most quantitatively important role in mitomycin C-enhanced Stx1 production by Stx phage H-19B [27], we found that integrase-deficient *C. rodentium* (Φ*stx_2dact_*), which is deficient in phage excision and replication (Table 4; [72]), produced levels of Stx2dact indistinguishable from wild type (Fig. 2). Thus, the ability to cause lethal Stx2-mediated disease was not correlated with the ability to generate infectious phage. Apparently, enhanced expression of late genes *stx_2dact_A* and *stx_2dac_B* still occurs in the absence of integrase and is sufficient for wild type levels of Stx2dact production. As expected, antiterminator protein Q, required for the transcription of late genes including *stx*, was essential for Stx2dact production by *C. rodentium* (Φ*stx_2dact_*), consistent with previous findings for the Stx2 phage Φ361 [26]. Finally, the S endolysin of Stx phage H-19B was previously shown to promote the timely release of toxin after mitomycin C induction [27]; we found that deficiency of the RS endolysins encoded by Φ*stx_2dact_* appeared to diminish the release of Stx2dact into culture supernatants at 4 hours post-induction (Fig. 2B). However, the decrease was not statistically significant, and RS-deficiency had no discernible effect on toxin release by 16 hours (Fig. 2C).

Whereas previous work in streptomycin-treated or gnotobiotic murine models has demonstrated that induction of the lytic developmental program of Stx phage occurs during infection and is required for disease [40, 41, 70], we document here that prophage induction occurs and is critical for productive infection of mice with intact microbiota. Our evidence includes first that *attP* sequences (indicative of excised, uningegrated phage genomes) were detected in the feces of infected mice, as revealed by deep sequencing (Table 3), or by qPCR (Fig. 3). Second, the ability of *C. rodentium* (Φ*stx_2dact_*) mutant derivatives to trigger the lytic cycle *in vitro* upon induction, with concomitant expression of the late *stx2_dact_* genes, correlated perfectly with the ability to cause lethal disease in mice. *C. rodentium* (Φ*stx_2dact_*) derivatives deficient in phage integrase, portal protein B, or host regulators RpoS or QseC were entirely competent for production of high levels of Stx2dact (Figs. 2 and S3), and upon infection of mice, each of these mutants retained the ability to cause weight loss and death, with kinetics indistinguishable from the wild type lysogen (Figs. 4 and S5). In contrast, prophage gene *Q*, critical for late gene transcription, was required for both *in vitro* Stx2 production and lethal infection. RecA, essential for the initiation of the SOS response that leads to prophage induction, was required for lethality after oral inoculation of *C. rodentium* (Φ*stx2dact*), consistent with the previous finding that RecA was required for lethality following intravenous EHEC infection of conventional mice [76]. Interestingly, human neutrophils and hydrogen peroxide are capable of increasing Stx production by EHEC *in vitro*, perhaps by triggering oxidative damage [77]. Our use of conventional C57BL/6 mice, for which many mutants are available, may facilitate studies to determine whether a specific inducing stimulus is responsible for lytic induction of *C. rodentium* (Φ*stx_2dact_*).

**Fig 4.**
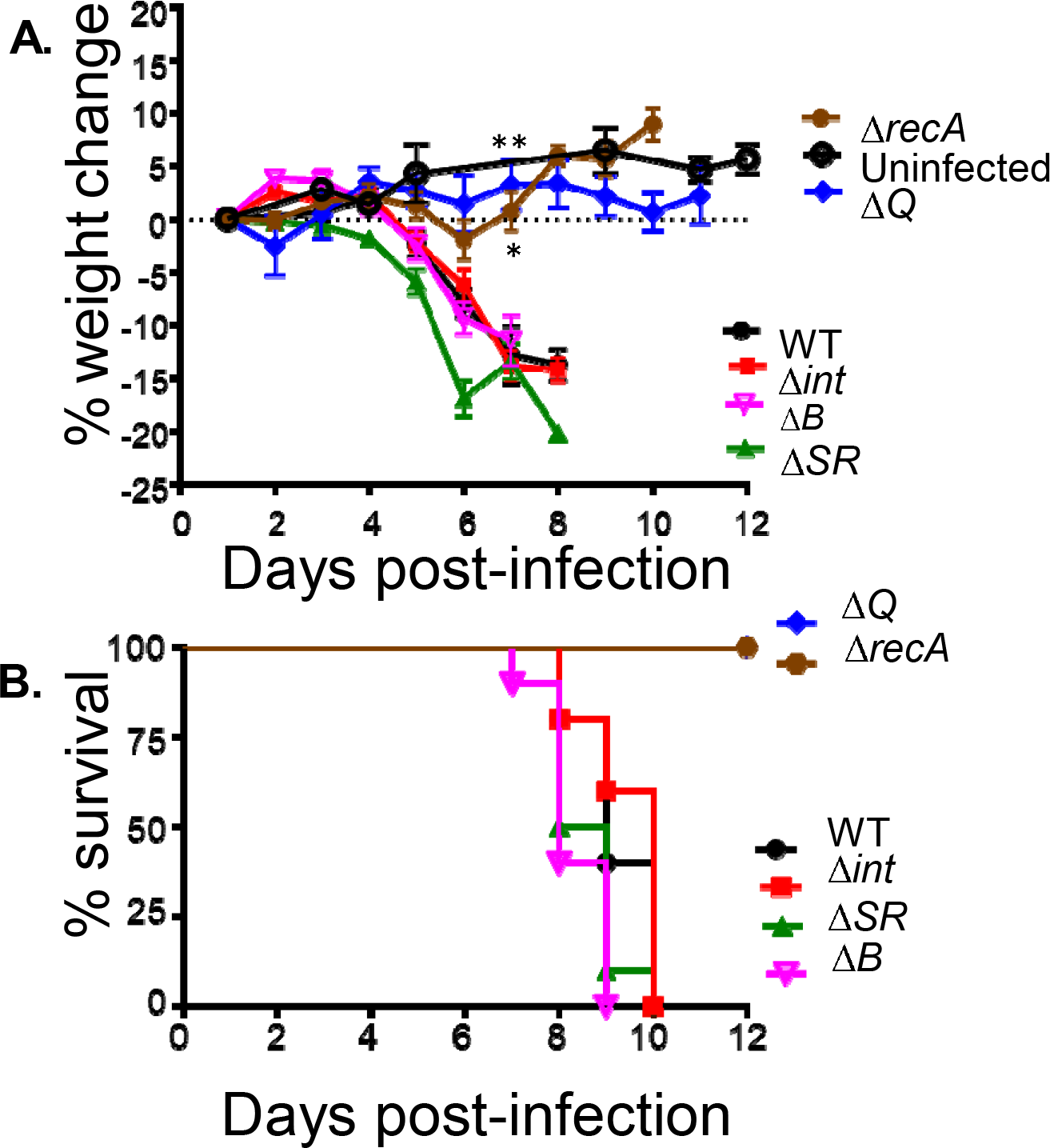
Lethal disease in mice correlates with the ability to produce Stx2dact but not with the ability to produce phage. Eight-week old female C57BL/6 mice were infected by oral gavage with the indicated lysogens. **A.** Percentage weight change was determined at indicated post-infection time. Data shown are averages ± SEM of 10 mice per group. Asterisks (*, **) indicate significance (p <0.05, <0.01) determined by 2-way ANOVA followed by Bonferroni post tests. **B.** Percent survival at the indicated post-infection time was monitored in 10 mice per group. Data represent cumulative results of 3 separate experiments.

We detected more than 1 ⨯ 10^9^ phage/ml in uninduced mid-log cultures, suggesting that there is a high level of spontaneous induction under *in vitro* culture conditions. In contrast, despite severe Stx2dact-mediated disease manifestations during productive infection by *C. rodentium* (Φ*stx_2dact_*), the number of *attP* sequences detected in feces was extremely low, suggesting that the level of prophage induction during infection may also be low. On day 6 post-infection, only 0.44% of all phage genomes detected were excised, compared to 99.66% that were integrated, reflecting intact prophage (Table 3). Depending on the day post-infection, excised phage detected by qPCR numbered 20- to 1000-fold fewer than viable *C. rodentium* (Φ*stx_2dact_*) cells (Fig. 3). Notably, previous work using a genetic reporter to indicate activation of lytic promoters of EHEC Stx phage 933W showed that the intestinal environment of a gnotobiotic mouse was strongly inducing [41]. While we cannot rule out the possibility that the low number of Φ*stx_2dact_ attP* sequences detected in feces reflects an instability of phage particles or some other factor in the intestinal milieu, our findings are consistent with the possibility that a low rate of Φ*stx_2dact_* induction may be sufficient to promote disease in this model, and that the presence or absence of the gut microbiota may have a consequential effect on disease outcome.

Consistent with the low level of phage detected in productively-infected mice, we found no evidence of Φ*stx_2dact_* lysogeny of commensal bacteria during *C. rodentium* (Φ*stx_dact_*) murine infection, suggesting that secondary infection of commensals by this phage is rare and that successive rounds of lytic infection are not an essential element of Stx production and disease in this microbiota-replete infection model. Given that the methods to measure phage particles utilized in this study can be applied to patient samples, future studies will focus on the extent of lytic induction of Stx phage during human infection, and how it may correlate with disease outcome.

## Materials and Methods

### Bacterial strains and plasmids

Strains and plasmids used in this study are listed in Table 1.

### Phage Φ*stx_2dact_* whole genome sequencing, assembly, and integration site determination

Genomic DNA was isolated from 5 ml of strain *C. rodentium* (Φ*stx_2dact_::kan^R^*) (Table 1) growr overnight at 37°C in LB broth containing chloramphenicol (12.5 μg/ml) and kanamycin (25 μg/ml). DNA was extracted using a DNeasy kit (Qiagen), according to the manufacturer’s protocol for Gram negative bacteria. A library of this DNA was then constructed for Illumina sequencing using Illumina TruSeq DNA Sample Preparation Kit per the manufacturer’s instructions. Following sequencing, the bacterial genome was assembled *de novo* into 1500 contigs using assemblers ABySS [57], and Edena [58]. The Bowtie2 program [59] was then used to map the *stx2* gene against this assembled genome and the contig containing this gene was identified. When aligned to the *C. rodentium* genome, a 69594-bp contig revealed a 47,343 bp prophage containing the *stx2* gene and other phage lambda-like gene sequences inserted into the host *dusA* gene. (Although the *C. rodentium dusA* gene is interrupted by the prophage genome, a potentially functional *dusA* gene is reconstituted at the *attL* bacterial/phage DNA junction by fusion with a prophage-derived open reading frame that we term “*ΦdusA’*” in Fig. 1.) The prophage sequence was deposited in GenBank as Φ1720a-02, accession number KF030445.1.

Integration of the prophage in both *C. rodentium (Φstx_2dact_*) and *C. rodentium* (ΦΔ*stx_2dact_::kan^R^*) into the host *dusA* gene was verified by PCR amplification of the *attL* and
*attR* phage-host junctions using primers DusF/PhageR and DusR/PhageF, respectively (Table 2), then DNA sequencing of the amplified junctions. Subsequent whole genome sequencing of *C. rodentium (Φstx_2dact_*) and *C. rodentium (ΦAstx_2dact_::kan^R^*) showed that, with the exception of the presence of the Φ*stx_2dact_ sequences*, they are identical to *C. rodentium* ICC 168, also known as strain DBS100 (GenBank accession number NC_013716.1), and to each other (data not shown).

### Phage Φ*stx_2dact_* genome annotation

The Φ*stx_2dact_* genome sequence was first annotated using the program RAST (http://rast.nmpdr.org/ [60]. The annotation was further refined by analyzing each open reading frame using the NCBI program MEGABLAST against the GenBank nucleotide database.

### Characterization of phage and prophage sequences in murine stool by massively parallel sequencing and analysis

DNA was extracted from fecal samples of 5 infected sick mice at 6 days post-infection, according to the method of Yang et al. [61]. Twenty mg stool samples were suspended in 5 ml PBS, pH7.2, and centrifuged at 100 ⨯ g for 15 min at 4°C. The supernatant was centrifuged at 13,000 ⨯ g for 10 min at 4°C, and the resulting pellet was washed 3 times in 1.5 ml acetone, centrifuging at 13,000 ⨯ g for 10 min at 4°C after each wash step. Two hundred μl of 5% Chelex-100 (Bio-Rad) and 0.2 mg proteinase K were added to the pellet and the sample was incubated for 30 min at 56°C. After vortexing briefly, the sample was centrifuged at 10,000 ⨯ g for 5 min and the supernatant containing the DNA was harvested and stored.

To characterize bacteria that harbor the Φ*stx_2dact_* prophage, we sequenced the bacterial bacterial-host *attL* prophage junction and adjacent bacterial DNA by following, with slight modifications (Suppl. Fig. 1), the methodology of Klein et al. [62] for constructing high-throughput sequencing libraries that contain a repetitive element (in this case, the phage *int* (integrase) gene). Briefly, genomic DNA was sheared by sonication to a size of 100-600 bp, followed by addition of ~20 deoxycytidine nucleotides to the 3’ ends of all molecules using Terminal deoxynucleotidyl Transferase (TdT, Fig. S1, step 2). Two rounds of PCR using a poly-C-specific and phage *int* gene-specific primer pair (PCR primers 1 and 2, Table 2) were used to amplify *attL* (Fig. S1, step 3) and to add on sequences necessary for high-throughput sequencing (PCR primers 3 and 4, Table 2, and Fig.S 1, step 4)).

Amplicons were sequenced using the MiSeq desktop sequencer (Ilumina) and primer Seq-P (Table 2), providing reads of up to 300 bp. As amplicons spanned the region from the phage *int* gene, through *attL*, and into the adjacent host genome (see Fig. 1B), reads of this length were required. 17,868,095 sequences encompassing 5 Gb were downloaded to the Galaxy server (https://usegalaxy.org/) and analyzed (Table 3). We first excluded sequences that clearly reflected *attL* (i.e., contained the 184 bp of Φ*dusA’*followed by *C. rodentium dusA*), indicating the prophage inserted into the *C. rodentium* genome. Of the remaining 801,959 sequences, 75,962 (0.44% of the total) encoded the intact *attP* site, implying that they were circular. These latter sequences presumably reflected excised circular phage genomes, possibly undergoing early theta DNA replication, ultimately leading to phage production. The remaining 725,997 sequences encoded only strings of A’s and/or C’s, and were eliminated from consideration.

### Generation of *C. rodentium* (Φ*stx*_2dact_) deletion constructs

Deletion mutants of *C. rodentium* (Φ*stx_2dact_*) in the prophage or the host genome were generated using a modified version of a one-step PCR-based gene inactivation protocol [63, 64]. Briefly, a PCR product of the zeocin-resistance gene and its promoter region flanked by 70-500 bp homology of the region upstream and downstream of the targeted gene was generated using the primers listed in Table 2. The chromosomal DNA served as template when the flanking regions were 500 bp in length on either side of the Zeocin cassette. The PCR product was electroporated into competent *C. rodentium* (Φ*stx_2dact_*) cells containing the lambda *red* plasmid pKD46 and recombinants were selected on plates containing chloramphenicol and zeocin (75 μg/ml). Replacement of the gene of interest with the zeocin resistance cassette was confirmed using specific primers (Table 2). At least two independent clones, validated using PCR, were obtained and subsequently analyzed.

### Quantification of Stx2 produced *in vitro*

Overnight 37°C cultures of *C. rodentium* (Φ*stx_2dact_*) or deletion derivatives were diluted 1:25 into 10 ml of fresh medium with appropriate antibiotics. Two independently derived clones for each mutant were tested, with indistinguishable results. The cultures were grown at 37°C with aeration to an OD_600_ of 0.4, and one ml of each culture was set aside (Table 4, “t=0h”) The remaining culture was split into 2 cultures. These cultures were grown for a further 4 hours (Table 4, “t=4h”) in the absence or presence of 0.25 μg/ml mitomycin C. (We first measured phage and Stx2 production at various times post-induction and found the 4-hour time point to be optimal for obtaining maximal phage and Stx2 following mitomycin C induction). Culture pellets and supernatants were then harvested by centrifugation at 17,800 ⨯ g for 5 minutes at room temperature. For *C. rodentium* (Φ*stx_2dact_*) and *C. rodentium* (Φ*stx_2dact_*Δ*SR*), a portion of each culture was also collected after ~16 h of incubation (“t=16h”). Supernatants and pellets were quantitated for Stx2 by ELISA, as described previously [52].

### Mouse infection studies

Mice were purchased from Jackson Laboratories and maintained in the Tufts University animal facility. All procedures were performed in compliance with Tufts University IACUC protocols. Seven to eight-week-old female C57BL/6J mice were gavaged with PBS or ~5⨯10^8^ CFU of overnight culture of *C. rodentium* (Φ*stx_2dact_*) or deletion derivatives in 100 μl PBS. Inoculum concentrations were confirmed by serial dilution plating. Fecal shedding was determined by plating dilutions of fecal slurry on either chloramphenicol, to detect wild type *C. rodentium* (Φ*stx_2dact_*), or chloramphenicol-zeocin plates, to detect deletion derivatives marked with a zeomycin resistance gene [52]. Body weights were monitored daily, and mice were euthanized upon losing >15% of their body weight.

DNA from infected mice fecal pellets was isolated using the QIAGEN DNeasy Blood and Tissue kit with modifications. Fecal pellets were incubated with buffer ATL and proteinase K overnight at 55°C. Buffer AL was added, and after mixing, pellets were further incubated at 56°C for 1 h. Pellet mixtures were then centrifuged at 8000 rpm for 1 min and the pellets were discarded. Ethanol was added to the supernatants, which were processed according to the manufacturer’s protocol. DNA concentrations were determined using a <u>NanoDrop™</u> spectrophotometer. qPCR was performed as described below.

### Quantification of phage genomes by qPCR

Excised phage genomes in cell supernatants were quantitated by qPCR. Supernatants were serially diluted 1:10, 1:100 and 1:1000 in distilled water. Separate reactions using two μl of the various dilutions as a template were carried out in duplicate. qPCR master-mix (Bio-Rad) was prepared according to the manufacturer’s instructions, using the *attP* primer set (Table 2) to detect copies of excised phage DNA. Results were compared to a standard curve, derived from a known concentration of a template fragments generated from amplifying *C. rodentium* (Φ*stx_2dac_*) DNA using *attP* primers. The template was serially diluted, in duplicate, to detect copy numbers ranging from 10^10^ to 10^2^. qPCR reactions were carried out as follows: 95°C for 3 min, followed by 35 cycles of 95°C for 1 min, 58°C for 30 sec, and 72°C for 1 min.

### Statistical tests

Data were analyzed using GraphPad Prism software. Comparison of multiple groups were performed using the Kruskal-Wallis test with Dunn’s multiple comparison post-test, or 2-way ANOVA with Bonferroni’s post-tests. In all tests, P values below 0.05 were considered statistically significant. Data represent the mean ± SEM in all graphs.

## Acknowledgements

We are grateful to Sara Roggensack for suggesting the PhageSeq methodology, David Lazinski for help with preparation of PhageSeq libraries, Martin Marinus for critical review of the manuscript, and Michael Pereira, Martin Marinus, and members of the Leong Lab for many useful suggestions.

## Supporting Information

**Fig S1.**
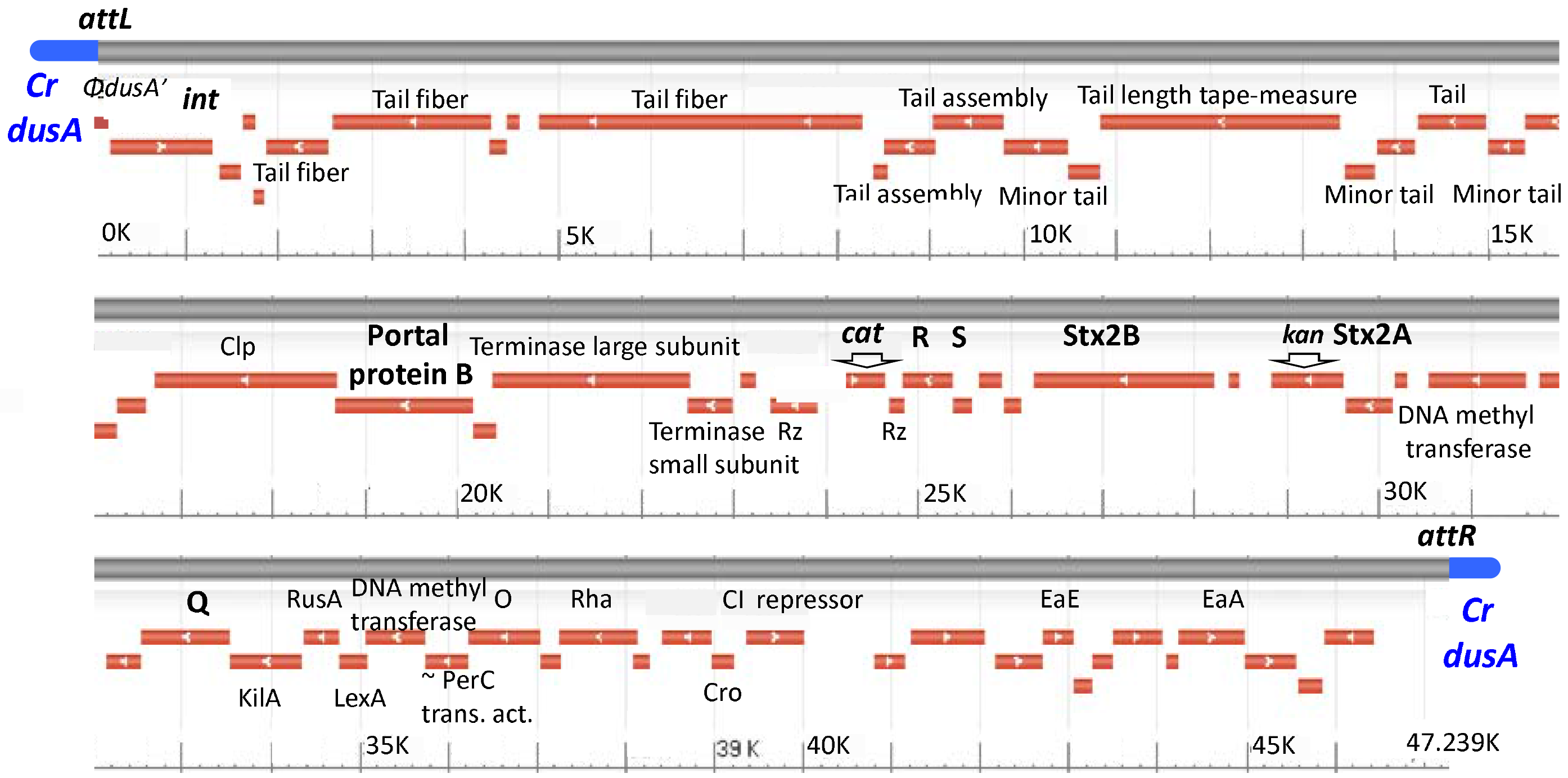
*C. rodentium* (Φ*stx_2dact_*) prophage annotation. The 47,239 bp prophage DNA sequence (gray), flanked by *attL* and *attR* upon insertion into *C. rodentium dusA* sequence (blue, “*Cr dusA*”), was determined by whole genome shotgun sequencing of *C. rodentium* (Φ*stx_2dact_,::kan^R^*) and annotated, as described in Materials and Methods. Names of encoded proteins are shown. Unannotated ORFs indicate hypothetical proteins. At the far left end is a phage sequence that encodes the N-terminal 112 amino acid of an open reading frame (“Φ*dusA*”) in the same reading frame as the 3’ end of the *C. rodentium dusA* gene. Strain *C. rodentium (Φstx_2dact_*) encodes a chloramphenicol acetyl transferase protein (*“cat”’*) inserted into the prophage *Rz* gene. The sequence of *C. rodentium* (Φ*stx2dact::kan^R^*) is identical to *C. rodentium* (Φ*stx_2dact_*) except that gene encoding the A subunit of Stx2dact (“Stx2A”) contains an 894 bp insertion encoding kanamycin resistance (“*kan*”), plus an additional 27 bp upstream and 28 bp downstream. Prophage genes studied in this work are shown in bold.
*Cr. C. rodentium*.

**Fig S2.**
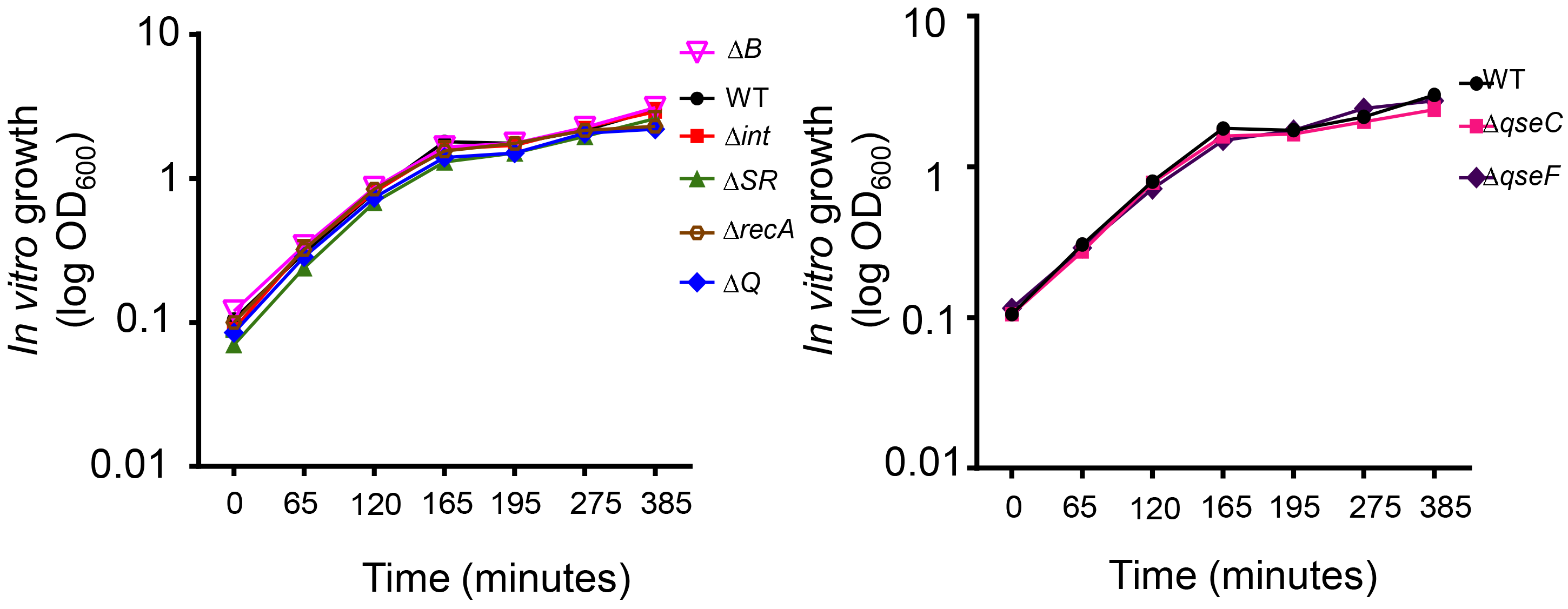
*C. rodentium (Φstx_2dact_*) mutants display no growth defects in rich medium. The indicated wild type or mutant *C. rodentium* (Φ*stx_2dact_act*) were grown in LB broth without antibiotics. Growth was measured over time by optical density (OD_600_), and growth curves are the average of duplicate samples. Doubling times were calculated based on the exponential growth regions of each curve. Representative results from one of two experiments are shown.

**Fig S3.**
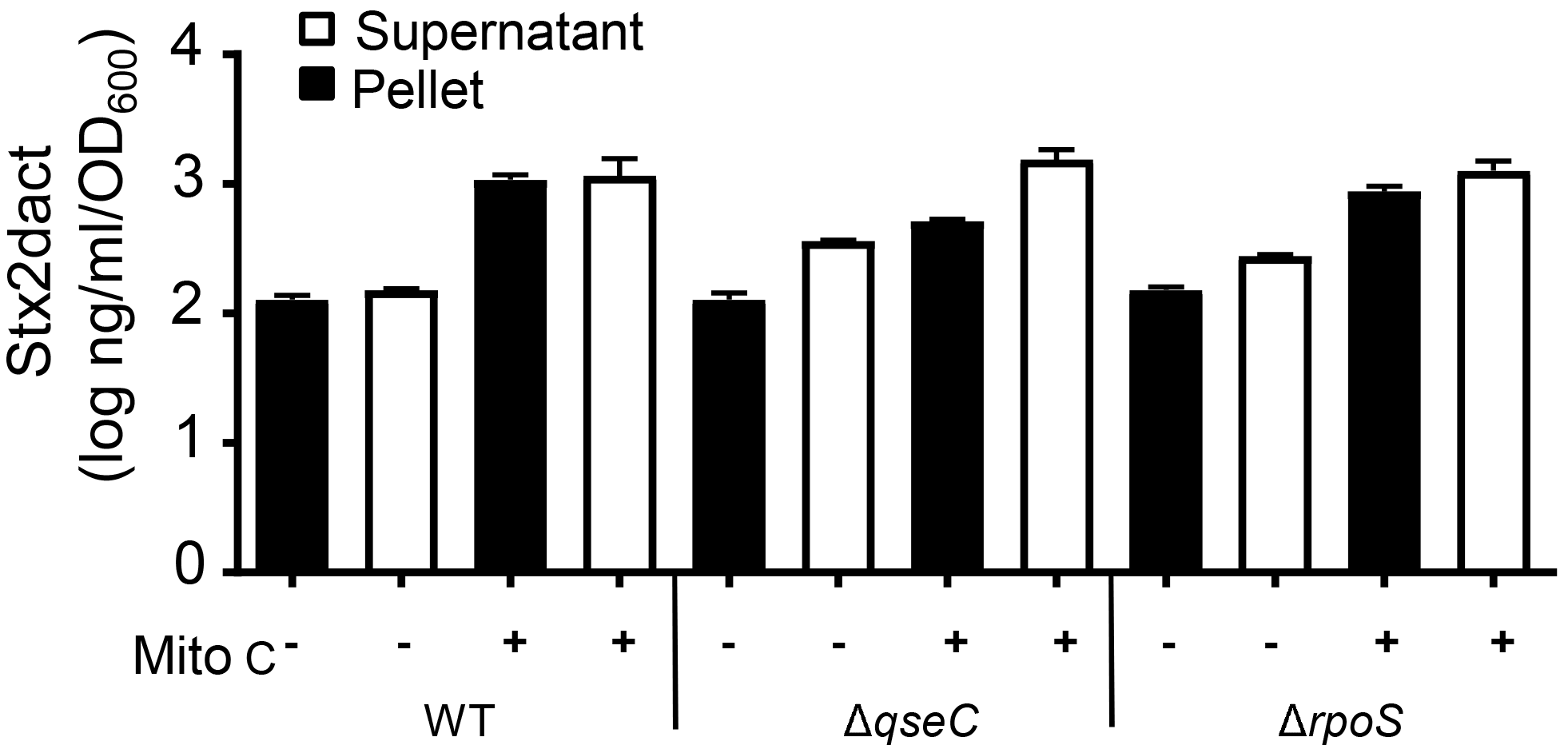
QseC and RpoS are not required for wild type basal and induced levels of Stx2 production *in vitro*. The indicated lysogens were grown to mid-log phase (designated as t=0h) and cultured for four more hours (t=4h) either in the absence (“-”) or presence (“+”) of 0.25 μg/ml mitomycin C. Pellets (filled bars) or supernatants (open bars) were subjected to capture ELISA to determine the level of Stx2 production. Quantities are expressed relative to the specific OD_600_ at t=0h. Results are averages ± SEM of triplicate samples, and are a representative of at least two experiments. Stx levels of neither the *C. rodentium* (Φ*stx_2dact_*Δ*qseC*) or *C. rodentium* (Φ*stx_2dact_* Δ*rpoS*) strains were significantly different from wild type *C. rodentium* (Φ*stx_2dact_act*), calculated using Kruskal-Wallis one-way analysis of variance followed by Dunn’s multiple comparisons test.

**Fig S4.**
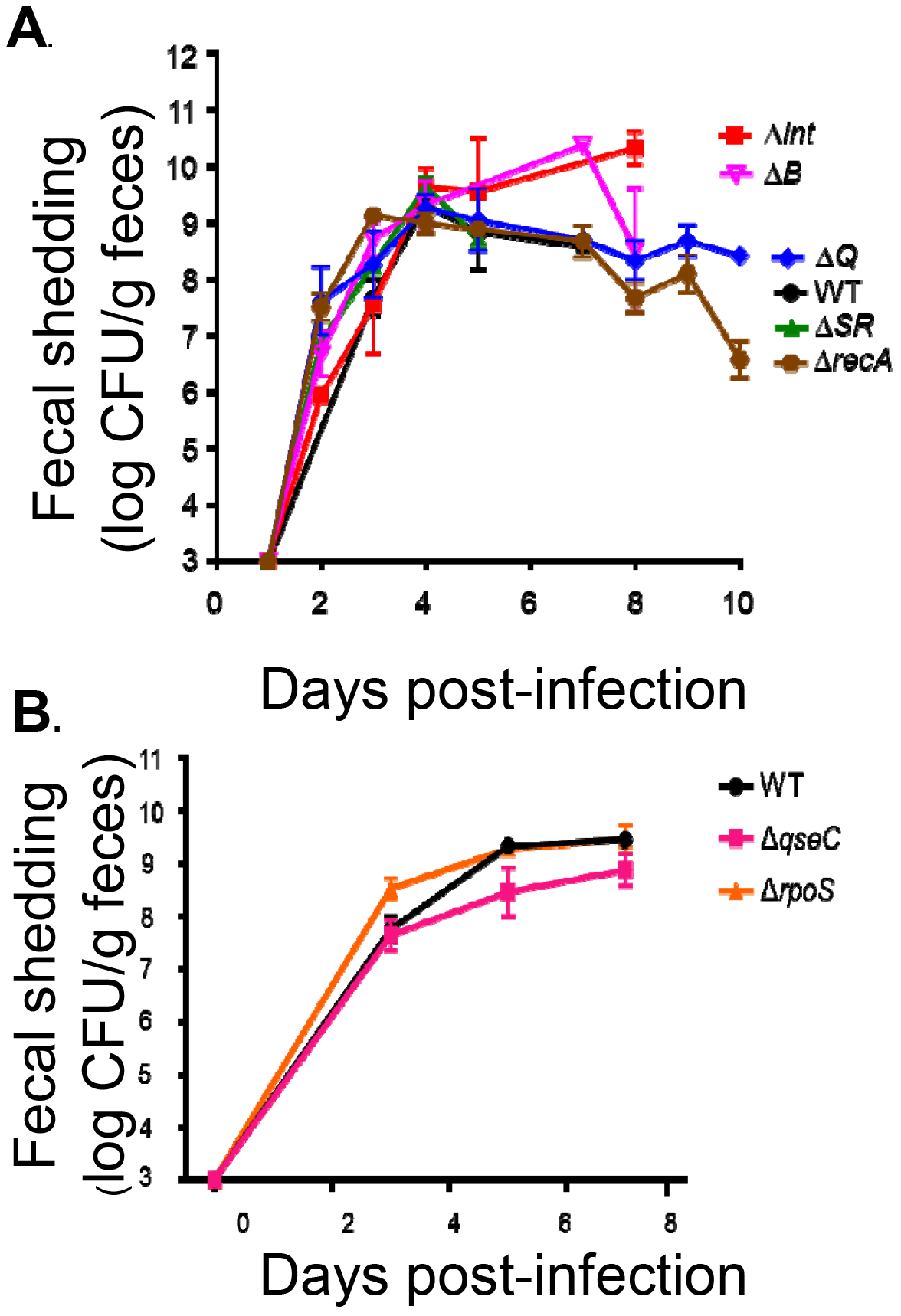
*C. rodentium* (Φ*stx_2dact_act*) mutants do not display colonization defects. Eight-week old female C57BL/6 mice were infected by oral gavage with the indicated lysogens. Fecal shedding of the lysogens was determined by plating for viable counts (see Materials and Methods). No significant differences were observed, as determined by 2-way ANOVA.

**Fig S5.**
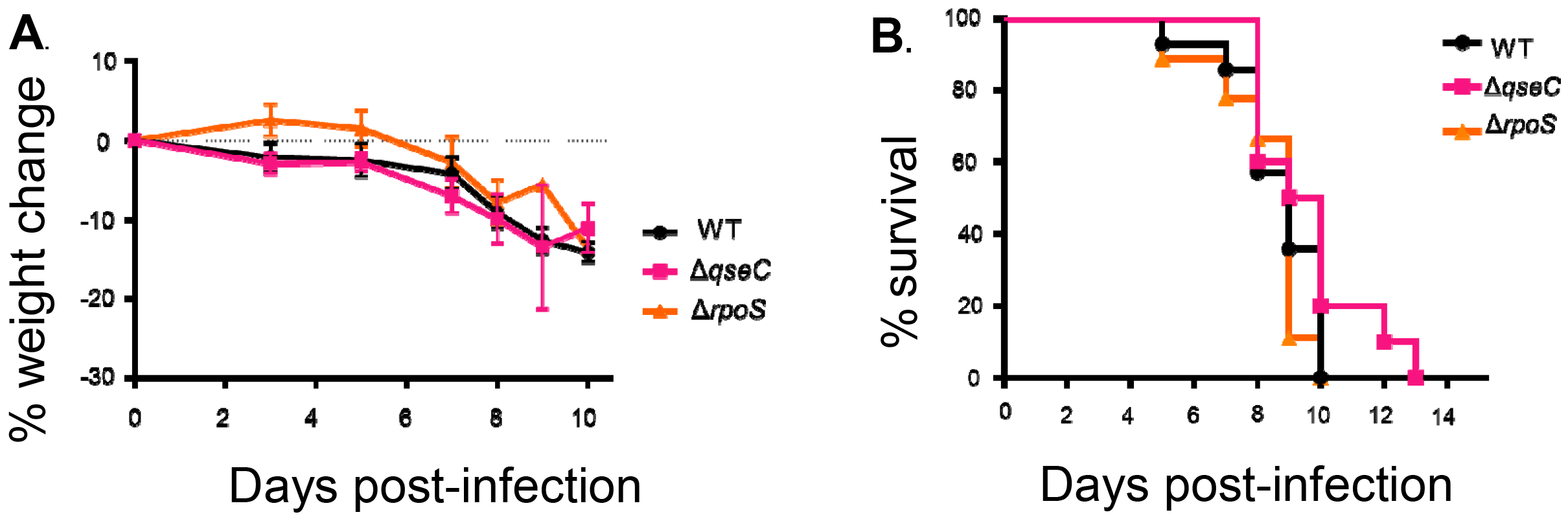
QseC and RpoS are not required for disease by *C. rodentium* (Φ*stx_2dact_act*). Eight-week old female C57BL/6 mice were infected by oral gavage with the indicated lysogens. **A.** Percentage weight change was determined at indicated post-infection time. Data shown are averages ± SEM of 10 mice per group. No significant differences were observed, as determined by 2-way ANOVA. **B.** Percent survival at the indicated post-infection time was monitored in 10 mice per group. Data represent cumulative results of 3 separate experiments.

## References

1. Karmali, M.A., V. Gannon, and J.M. Sargeant, Verocytotoxin-producing Escherichia coli (VTEC). Vet Microbiol, 2010. 140(3-4): p. 360–70.

2. Kaper, J.B., J.P. Nataro, and H.L. Mobley, Pathogenic Escherichia coli. Nat Rev Microbiol, 2004. 2(2): p. 123–40.

3. Pennington, H., Escherichia coli O157. Lancet, 2010. 376(9750): p. 1428–35.

4. Sperandio, V. and C.J. Hovde, eds. Enterohemorrhagic Escherichia coli and Other Shiga Toxin-Producing E. coli.. 2015, ASM Press: Washington, D.C.

5. Karmali, M.A., Host and pathogen determinants of verocytotoxin-producing Escherichia coli-associated hemolytic uremic syndrome. Kidney Int Suppl, 2009(112): p. S4–7.

6. Tarr, P.I., C.A. Gordon, and W.L. Chandler, Shiga-toxin-producing Escherichia coli and haemolytic uraemic syndrome. Lancet, 2005. 365(9464): p. 1073–86.

7. Scheiring, J., et al., Outcome in patients with recurrent hemolytic uremic syndrome. Pediatric Transplantation, 2005. 9: p. 48–48.

8. Brady, M.J., et al., Enterohaemorrhagic and enteropathogenic Escherichia coli Tir proteins trigger a common Nck-independent actin assembly pathway. Cell Microbiol, 2007. 9(9): p. 2242–53.

9. Vingadassalom, D., et al., Insulin receptor tyrosine kinase substrate links the E. coli O157:H7 actin assembly effectors Tir and EspF(U) during pedestal formation. Proc Natl Acad Sci U S A, 2009. 106(16): p. 6754–9.

10. Lai, Y., et al., Intimate host attachment: enteropathogenic and enterohaemorrhagic Escherichia coli. Cell Microbiol, 2013. 15(11): p. 1796–808.

11. Proulx, F., E.G. Seidman, and D. Karpman, Pathogenesis of Shiga toxin-associated hemolytic uremic syndrome. Pediatr Res, 2001. 50(2): p. 163–71.

12. Thorpe, C.M. and D.W. Acheson, Testing of urinary Escherichia coli isolates for Shiga toxin production. Clin Infect Dis, 2001. 32(10): p. 1517–8.

13. Robinson, C.M., et al., Shiga toxin of enterohemorrhagic Escherichia coli type O157:H7 promotes intestinal colonization. Proc Natl Acad Sci U S A, 2006. 103(25): p. 9667–72.

14. Obrig, T.G., Escherichia coli Shiga Toxin Mechanisms of Action in Renal Disease. Toxins (Basel), 2010. 2(12): p. 2769–2794.

15. Melton-Celsa, A., et al., Pathogenesis of Shiga-toxin producing escherichia coli. Curr Top Microbiol Immunol, 2012. 357: p. 67–103.

16. Wadolkowski, E.A., J.A. Burris, and A.D. O’Brien, Mouse model for colonization and disease caused by enterohemorrhagic Escherichia coli O157:H7. Infect Immun, 1990. 58(8): p. 2438–45.

17. Wadolkowski, E.A., et al., Acute renal tubular necrosis and death of mice orally infected with Escherichia coli strains that produce Shiga-like toxin type II. Infect Immun, 1990. 58(12): p. 3959–65.

18. Keepers, T.R., et al., A murine model of HUS: Shiga toxin with lipopolysaccharide mimics the renal damage and physiologic response of human disease. J Am Soc Nephrol, 2006. 17(12): p. 3404–14.

19. Davis, T.K., N.C. Van De Kar, and P.I. Tarr, Shiga Toxin/Verocytotoxin-Producing Escherichia coli Infections: Practical Clinical Perspectives. Microbiol Spectr, 2014. 2(4): p. EHEC-0025–2014.

20. Melton-Celsa, A.R. and A.D. O’Brien, New Therapeutic Developments against Shiga Toxin-Producing Escherichia coli. Microbiol Spectr, 2014. 2(5).

21. Freedman, S.B., et al., Shiga Toxin-Producing Escherichia coli Infection, Antibiotics, and Risk of Developing Hemolytic Uremic Syndrome: A Meta-analysis. Clin Infect Dis, 2016. 62(10): p. 1251–1258.

22. Mizutani, S., N. Nakazono, and Y. Sugino, The so-called chromosomal verotoxin genes are actually carried by defective prophages. DNA Res, 1999. 6(2): p. 141–3.

23. Tyler, J.S., J. Livny, and D.I. Friedman, Lambdoid Phages and Shiga Toxin., in Phages; Their role in Pathogenesis and Biotechnology., M.K. Waldor, D.I. Friedman, and S. Adhya, Editors. 2005, ASM Press: Washington, D.C. p. 131–164.

24. Neely, M.N. and D.I. Friedman, Arrangement and functional identification of genes in the regulatory region of lambdoid phage H-19B, a carrier of a Shiga-like toxin. Gene, 1998. 223(1-2): p. 105–13.

25. Neely, M.N. and D.I. Friedman, Functional and genetic analysis of regulatory regions of coliphage H-19B: location of shiga-like toxin and lysis genes suggest a role for phage functions in toxin release. Mol Microbiol, 1998. 28(6): p. 1255–67.

26. Wagner, P.L., et al., Role for a phage promoter in Shiga toxin 2 expression from a pathogenic Escherichia coli strain. J Bacteriol, 2001. 183(6): p. 2081–5.

27. Wagner, P.L., et al., Bacteriophage control of Shiga toxin 1 production and release by Escherichia coli. Mol Microbiol, 2002. 44(4): p. 957–70.

28. Tyler, J.S., M.J. Mills, and D.I. Friedman, The operator and early promoter region of the Shiga toxin type 2-encoding bacteriophage 933W and control of toxin expression. J Bacteriol, 2004. 186(22): p. 7670–9.

29. Casjens, S.R. and R.W. Hendrix, Bacteriophage lambda: Early pioneer and still relevant. Virology, 2015. 479-480: p. 310–30.

30. Hughes, D.T., et al., The QseC adrenergic signaling cascade in Enterohemorrhagic E. coli (EHEC). PLoS Pathog, 2009. 5(8): p. e1000553.

31. Imamovic, L., et al., Heterogeneity in phage induction enables the survival of the lysogenic population. Environ Microbiol, 2016. 18(3): p. 957–69.

32. Karch, H., N.A. Strockbine, and A.D. O’Brien, Growth of Escherichia coli in the presence of trimethoprim-sulfamethoxazole facilitates detection of Shiga-like toxin producing strains by colony blot assay FEMS Microbiol Lett, 1986. 35(2-3): p. 141145.

33. Walterspiel, J.N., et al., Effect of subinhibitory concentrations of antibiotics on extracellular Shiga-like toxin I. Infection, 1992. 20(1): p. 25–9.

34. Matsushiro, A., et al., Induction of prophages of enterohemorrhagic Escherichia coli O157:H7 with norfloxacin. J Bacteriol, 1999. 181(7): p. 2257–60.

35. Wong, C.S., et al., The Risk of the Hemolytic-Uremic Syndrome after Antibiotic Treatment of Escherichia coli O157:H7 Infections. The New England Journal of Medicine, 2000. 342: p. 1930–1936.

36. Dundas, S., et al., Using antibiotics in suspected haemolytic-uraemic syndrome: antibiotics should not be used in Escherichia coli O157:H7 infection. BMJ, 2005. 330(7501): p. 1209; author reply 1209.

37. Mohawk, K.L., et al., Pathogenesis of Escherichia coli O157:H7 strain 86-24 following oral infection of BALB/c mice with an intact commensal flora. Microb Pathog, 2010. 48(3-4): p. 131–42.

38. Goswami, K., et al., Coculture of Escherichia coli O157:H7 with a Nonpathogenic E. coli Strain Increases Toxin Production and Virulence in a Germfree Mouse Model. Infect Immun, 2015. 83(11): p. 4185–93.

39. Taguchi, H., et al., Experimental infection of germ-free mice with hyper-toxigenic enterohaemorrhagic Escherichia coli O157:H7, strain 6. J Med Microbiol, 2002. 51(4): p. 336–43.

40. Zhang, X., et al., Quinolone antibiotics induce Shiga toxin-encoding bacteriophages, toxin production, and death in mice. J Infect Dis, 2000. 181(2): p. 664–70.

41. Tyler, J.S., et al., Prophage induction is enhanced and required for renal disease and lethality in an EHEC mouse model. PLoS Pathog, 2013. 9(3): p. e1003236.

42. Fiebiger, U., S. Bereswill, and M.M. Heimesaat, Dissecting the Interplay Between Intestinal Microbiota and Host Immunity in Health and Disease: Lessons Learned from Germfree and Gnotobiotic Animal Models. Eur J Microbiol Immunol (Bp), 2016. 6(4): p. 253–271.

43. Myhal, M.L., D.C. Laux, and P.S. Cohen, Relative colonizing abilities of human fecal and K 12 strains of Escherichia coli in the large intestines of streptomycin-treated mice. Eur J Clin Microbiol, 1982. 1(3): p. 186–92.

44. Gamage, S.D., et al., Commensal bacteria influence Escherichia coli O157:H7 persistence and Shiga toxin production in the mouse intestine. Infect Immun, 2006. 74(3): p. 1977–83.

45. Gamage, S.D., et al., Nonpathogenic Escherichia coli can contribute to the production of Shiga toxin. Infect Immun, 2003. 71(6): p. 3107–15.

46. Gamage, S.D., et al., Diversity and host range of Shiga toxin-encoding phage. Infect Immun, 2004. 72(12): p. 7131–9.

47. Iversen, H., et al., Commensal E. coli Stx2 lysogens produce high levels of phages after spontaneous prophage induction. Front Cell Infect Microbiol, 2015. 5: p. 5.

48. de Sablet, T., et al., Human microbiota-secreted factors inhibit shiga toxin synthesis by enterohemorrhagic Escherichia coli O157:H7. Infect Immun, 2009. 77(2): p. 78390.

49. Asahara, T., et al., Probiotic bifidobacteria protect mice from lethal infection with Shiga toxin-producing Escherichia coli O157:H7. Infect Immun, 2004. 72(4): p. 2240–7.

50. Carey, C.M., et al., The effect of probiotics and organic acids on Shiga-toxin 2 gene expression in enterohemorrhagic Escherichia coli O157:H7. J Microbiol Methods, 2008. 73(2): p. 125–32.

51. Toshima, H., et al., Enhancement of Shiga toxin production in enterohemorrhagic Escherichia coli serotype O157:H7 by DNase colicins. Appl Environ Microbiol, 2007. 73(23): p. 7582–8.

52. Mallick, E.M., et al., A novel murine infection model for Shiga toxin-producing Escherichia coli. J Clin Invest, 2012. 122(11): p. 4012–24.

53. Mallick, E.M., et al., The ability of an attaching and effacing pathogen to trigger localized actin assembly contributes to virulence by promoting mucosal attachment. Cell Microbiol, 2014. 16(9): p. 1405–24.

54. Gobius, K.S., G.M. Higgs, and P.M. Desmarchelier, Presence of activatable Shiga toxin genotype (stx(2d)) in Shiga toxigenic Escherichia coli from livestock sources. J Clin Microbiol, 2003. 41(8): p. 3777–83.

55. Melton-Celsa, A.R., S.C. Darnell, and A.D. O’Brien, Activation of Shiga-like toxins by mouse and human intestinal mucus correlates with virulence of enterohemorrhagic Escherichia coli O91:H21 isolates in orally infected, streptomycin-treated mice. Infect Immun, 1996. 64(5): p. 1569–76.

56. Teel, L.D., et al., One of two copies of the gene for the activatable shiga toxin type 2d in Escherichia coli O91:H21 strain B2F1 is associated with an inducible bacteriophage. Infect Immun, 2002. 70(8): p. 4282–91.

57. Simpson, J.T., et al., ABySS: A parallel assembler for short read sequence data. Genome Research, 2009. 19(6): p. 1117–1123.

58. Hernandez, D., et al., De novo bacterial genome sequencing: Millions of very short reads assembled on a desktop computer. Genome Research, 2008. 18(5): p. 802809.

59. Langmead, B. and S.L. Salzberg, Fast gapped-read alignment with Bowtie 2. Nat Methods, 2012. 9(4): p. 357–9.

60. Aziz, R.K., et al., The RAST Server: Rapid Annotations using Subsystems Technology. BMC Genomics 2008. 9: p. 75.

61. Yang, J.-L., et al., A simple and rapid method for extracting bacterial DNA from intestinal microflora for ERIC-PCR detection. World J. Gastroenterol., 2008. 14: p. 2872–2876.

62. Klein, B.A., et al., Identification of essential genes of the periodontal pathogen Porphyromonas gingivalis. BMC Genomics, 2012. 13: p. 578.

63. Datsenko, K.A. and B.L. Wanner, One-step inactivation of chromosomal genes in Escherichia coli K-12 using PCR products. Proc Natl Acad Sci U S A, 2000. 97(12): p. 6640–5.

64. Mallick, E.M., et al., Allele-and tir-independent functions of intimin in diverse animal infection models. Front Microbiol, 2012. 3: p. 11.

65. Smith, D.L., et al., Comparative genomics of Shiga toxin encoding bacteriophages. BMC Genomics, 2012. 13: p. 311.

66. Gottesman, M.E. and R.A. Weisberg, Little lambda, who made thee? Microbiol Mol Biol Rev, 2004. 68(4): p. 796–813.

67. Farrugia, D.N., et al., A novel family of integrases associated with prophages and genomic islands integrated within the tRNA-dihydrouridine synthase A (dusA) gene. Nucleic Acids Res, 2015. 43(9): p. 4547–57.

68. Nash, H.A., Integration and excision of bacteriophage lambda: the mechanism of conservation site specific recombination. Annu Rev Genet, 1981. 15: p. 143–67.

69. Petty, N.K., et al., The Citrobacter rodentium genome sequence reveals convergent evolution with human pathogenic Escherichia coli. J Bacteriol, 2010. 192(2): p. 525–38.

70. Acheson, D.W., et al., In vivo transduction with shiga toxin 1-encoding phage. Infect Immun, 1998. 66(9): p. 4496–8.

71. Anderson, B., et al., Enumeration of bacteriophage particles: Comparative analysis of the traditional plaque assay and real-time QPCR-and nanosight-based assays. Bacteriophage, 2011. 1(2): p. 86–93.

72. Stevens, W.F., S. Adhya, and W. Szybalski, Origin and bidirectional orientation of DNA Replication in coliphage lambda in The Bacteriophage Lambda., A.D. Hershey, Editor. 1971, Cold Spring Harbor Laboratory: Cold Spring Harbor, NY. p. 515–533.

73. Gottesman, M.E. and M.B. Yarmolinsky, Integration-Negative Mutants of Bacteriophage Lambda. Journal of Molecular Biology, 1968. 31(3): p. 487-+.

74. Dove, W.F., Action of the lambda chromosome. I. Control of functions late in bacteriophage development. J Mol Biol, 1966. 19(1): p. 187–201.

75. Moreira, C.G., et al., Bacterial Adrenergic Sensors Regulate Virulence of Enteric Pathogens in the Gut. MBio, 2016. 7(3).

76. Fuchs, S., et al., Influence of RecA on in vivo virulence and Shiga toxin 2 production in Escherichia coli pathogens. Microb Pathog, 1999. 27(1): p. 13–23.

77. Wagner, P.L., D.W. Acheson, and M.K. Waldor, Human neutrophils and their products induce Shiga toxin production by enterohemorrhagic Escherichia coli. Infect Immun, 2001. 69(3): p. 1934–7.

78. Mallick, E.M., et al., A novel murine infection model for Shiga toxin-producing Escherichia coli. The Journal of clinical investigation, 2012.

